# The plastid genome of the non-photosynthetic plant *Rhopalocnemis phalloides* is one of the most polymorphic genomes known

**DOI:** 10.1101/2025.10.06.680833

**Authors:** Mikhail I. Schelkunov, Maxim S. Nuraliev, Maria D. Logacheva

## Abstract

The common ancestor of plants in the Balanophoraceae family lost the ability to photosynthesize, and all several dozen extant species of this family feed exclusively by parasitizing other plants. Previous studies have shown that the plastid genomes of Balanophoraceae present numerous unusual features, including an extraordinarily increased rate of mutation accumulation without an observable weakening of natural selection acting on the genes.

This study aims to test the hypothesis that the increased rate of mutation accumulation could have led to exceptionally high levels of intraspecific polymorphism in Balanophoraceae. To this end, we studied the plastid genomes of 7 samples of the species *Rhopalocnemis phalloides*.

Although all 7 plastid genomes possessed the same genes arranged in the same order, the level of polymorphism was indeed extremely high, possibly the highest among all known genomes of any living organisms. Specifically, the average genome-wide percent identity of the plastid genomes of *Rhopalocnemis phalloides* was 68.7%, and the average percent identity of the CDSs was 68.9%.

Additionally, during this study, we discovered 60 taxonomically misclassified plastid genomes in NCBI databases; this result has independent value.

## Introduction

Balanophoraceae is a family of flowering plants consisting of several dozen species (Hansen 2015). The exact number is still unknown because scientists continue to discover new species in this family (Yu et al. 2021; Cardoso & Braga 2024). An unusual characteristic of Balanophoraceae is that all members of this family lack the ability to photosynthesize and, instead, parasitize other plants (Hansen 2015). However, the ancestors of Balanophoraceae were typical photosynthetic flowering plants.

The transition to a non-photosynthetic lifestyle led to numerous genomic changes in Balanophoraceae (Sanchez-Puerta et al. 2017; Schelkunov et al. 2019; Su et al. 2019; Schelkunov et al. 2021; Roulet et al. 2020; Ceriotti et al. 2021; Yu et al. 2022; Chen et al. 2023; Kim et al. 2023; Zhou et al. 2023; Ceriotti et al. 2025; Svetlikova et al. 2025). In particular, they experienced a massive loss of genes related to photosynthesis in both the plastid and nuclear genomes. Additionally, the AT content in the plastid genomes of Balanophoraceae is extremely high, approximately 80–90% in most of the studied species. Additionally, the rate of mutation accumulation in the plastid genomes of Balanophoraceae is 1–2 orders of magnitude higher than that in photosynthetic plants. The increase in AT content and mutation accumulation rate is apparently explained by the loss of some nuclear-encoded genes responsible for plastid genome repair (Schelkunov et al. 2021; Ceriotti et al. 2022; Chen et al. 2023). The reason for the loss of these genes is unknown, although there are hypotheses (Schelkunov et al. 2021).

Previous studies have shown that among the members of the Balanophoraceae family with known plastid genomes, *Rhopalocnemis phalloides* (Figure 1a) has one of the highest mutation accumulation rates in the plastid genome (Ceriotti et al. 2025; Kim et al. 2023; Svetlikova et al. 2025). Since *Rhopalocnemis phalloides* is apparently the only species of its genus, we refer to it simply as “rhopalocnemis”. The main objective of this study was to test the hypothesis that due to the high rate of mutation accumulation, the plastid genome of rhopalocnemis has become extremely polymorphic. Indeed, in this article, we show that it is the most polymorphic among all known plastid genomes of flowering plants and, moreover, that it may be the most polymorphic of all known genomes.

**Figure 1.**
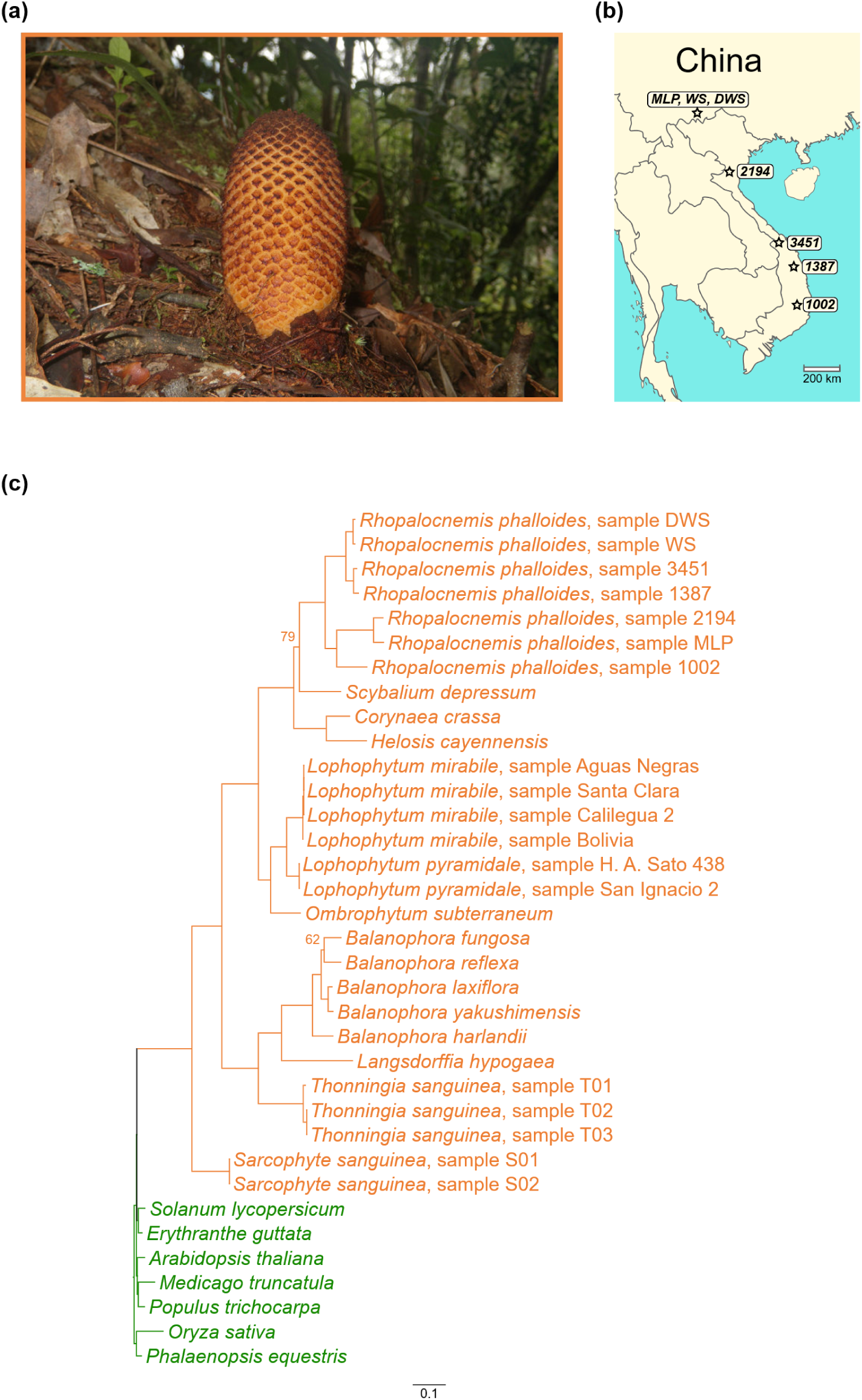
(a) The inflorescence of rhopalocnemis—the only part of rhopalocnemis visible above ground. (b) Map with sample collection sites marked with stars. Three samples were collected from nearby locations in China, and four samples were collected in Vietnam. (c) Phylogenetic tree of the studied species. Species of the Balanophoraceae are marked in orange, and photosynthetic species are marked in green. UFBoot support values equal to 100 are not labeled. The line at the bottom indicates the number of substitutions per site. The tree shows an increased rate of mutation accumulation in Balanophoraceae, especially in rhopalocnemis, compared to photosynthetic plants.

## Results and Discussion

### On the metrics of genetic polymorphism used in this study

There are many metrics of genetic polymorphism. In this study, we use three of them (Table 1). Each of these metrics, as shown in the table, has its own advantages and disadvantages. As the main metric of genetic polymorphism in this article, we use MPPI because this metric, in our opinion, is the most intuitive. The values of the other two metrics are provided mainly in the Supplementary section.

**Table 1.**
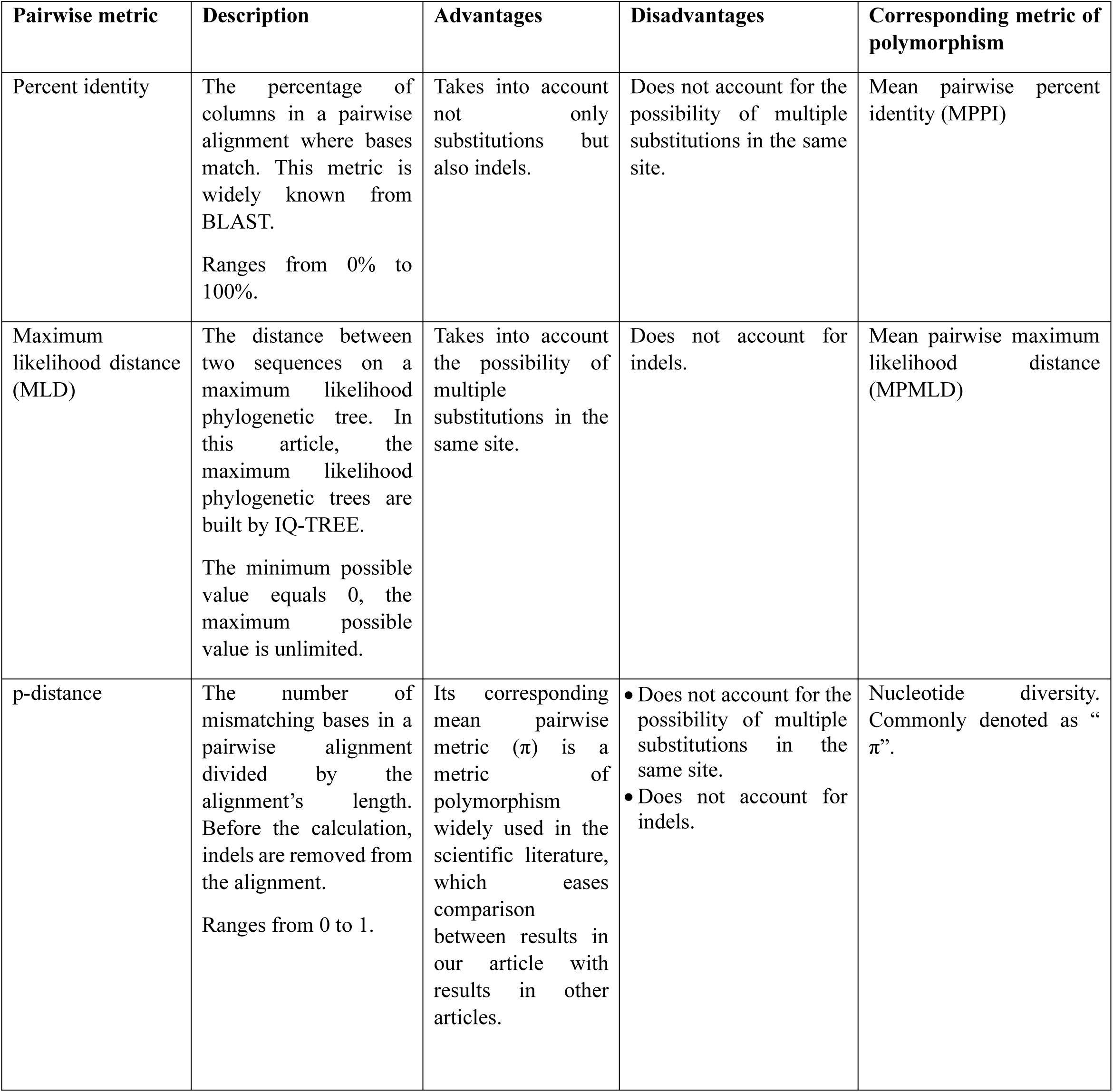
Metrics of sequence similarity and dissimilarity used in this article.

| Pairwise metric | Description | Advantages | Disadvantages | Corresponding metric of polymorphism |
| --- | --- | --- | --- | --- |
| Percent identity | The percentage of columns in a pairwise alignment where bases match. This metric is widely known from BLAST.<br>Ranges from 0% to 100%. | Takes into account not only substitutions but also indels. | Does not account for the possibility of multiple substitutions in the same site. | Mean pairwise percent identity (MPPI) |
| Maximum likelihood distance (MLD) | The distance between two sequences on a maximum likelihood phylogenetic tree. In this article, the maximum likelihood phylogenetic trees are built by IQ-TREE.<br>The minimum possible value equals 0, the maximum possible value is unlimited. | Takes into account the possibility of multiple substitutions in the same site. | Does not account for indels. | Mean pairwise maximum likelihood distance (MPMLD) |
| p-distance | The number of mismatching bases in a pairwise alignment divided by the alignment's length. Before the calculation, indels are removed from the alignment.<br>Ranges from 0 to 1. | Its corresponding mean pairwise metric ($\pi$) is a metric of polymorphism widely used in the scientific literature, which eases comparison between results in our article with results in other articles. | • Does not account for the possibility of multiple substitutions in the same site.<br>• Does not account for indels. | Nucleotide diversity. Commonly denoted as "$\pi$". |

MPPI is the arithmetic mean of pairwise percent identities between sequences. Just as the percent identity between sequences can range from 0% to 100%, MPPI can also range from 0% to 100%. For details on the calculation of the percent identity, MPPI, and other metrics of genetic similarity and dissimilarity, see the Materials and Methods section.

MPPI, MPMLD, and π are correlated with each other. The p-value for each of the three pairwise correlations, calculated using t-tests with a Bonferroni correction, is less than 10^-9^ (Supplementary Figure 1).

### Basic characteristics of the plastid genomes of rhopalocnemis

In this study, we examined the plastid genomes of 7 samples of rhopalocnemis (Figure 1b). Three of them were sequenced by us specifically for this study, one was previously sequenced by us, and three were sequenced by other scientists. It should be noted that in the assemblies of two plastid genomes of rhopalocnemis available in the NCBI databases, we found minor errors and corrected them (see Supplementary Note 1). All analyses in this article are based on corrected versions of the genomes.

The main features of the plastid genomes of the 7 samples of rhopalocnemis are presented in Table 2. In the same table, we have included the features of 21 plastid genomes of other species of Balanophoraceae and, for comparison, of 7 plastid genomes from model species of photosynthetic flowering plants.

The plastid genomes of all 7 samples of rhopalocnemis have the same 16 genes, and the order of these genes is the same. In other words, all 7 plastid genomes are collinear. These 16 genes are as follows:

1. 9 genes encoding protein components of the plastid ribosome: *rps3, rps7, rps12, rps14, rps18, rps19, rpl2, rpl16, rpl36*;
2. 4 genes encoding proteins with other functions: *accD, clpP, ycf1, ycf2*;
3. 2 genes encoding RNA components of the plastid ribosome: *rrn16, rrn23*;
4. 1 gene that previously encoded a tRNA for glutamic acid but now apparently has a different function. This gene is discussed in a separate section titled “What happened to *trnE*?”.

A description of the features of these genes, such as the average length, dN/dS, and other features, can be found in Supplementary Table 1.

A detailed description of the plastid genome of rhopalocnemis has already been provided in the article of Schelkunov et al. (2019), so we will not dwell on it here. However, in that article, we considered the *trnE* gene a pseudogene, whereas we have now changed our perspective on this issue.

In the plastid genomes of several genera of the Balanophoraceae family, a change in the genetic code has previously been observed. Specifically, in *Ombrophytum* and *Lophophytum*, TGA ceased to be a stop codon and became a tryptophan codon (Ceriotti et al. 2021). In *Balanophora*, TAG ceased to be a stop codon and became a tryptophan codon (Su et al. 2019). In all 7 samples of rhopalocnemis, we observe the standard genetic code, although in some samples (simply due to the short genome length), some GC-rich codons do not occur (Supplementary Table 2). For example, the alanine codon GCG occurs once in samples MLP and WS but does not occur at all in the other five samples.

As shown in Table 2, all samples of Balanophoraceae have reduced plastid genomes compared to photosynthetic plants; this reduction is associated with the loss of genes directly or indirectly related to photosynthesis. Additionally, the AT content has increased in Balanophoraceae. Both of these characteristics, typical not only of Balanophoraceae but also of other lineages of flowering plants that have lost the ability to photosynthesize (for example, some orchids and broomrapes), have been extensively described in the literature (Barbrook et al. 2006; Graham et al. 2017; Hadariová et al. 2018; Wicke & Naumann 2018).

The phylogenetic tree of the studied plastid genomes is shown in Figure 1c. Compared with that in photosynthetic plants, the rate of mutation accumulation is considerably increased in rhopalocnemis and other members of the Balanophoraceae.

**Table 2.** The plastid genomes used in this article and their basic features. The first 7 plastid genomes belong to model photosynthetic plants; the remainder are from members of the Balanophoraceae, all of which are non-photosynthetic. Duplicated genes with identical sequences were counted as a single gene. ^a^ - This sample of *Lophophytum pyramidale* is deposited on the NCBI website under a synonymous (according to the World Checklist of Vascular Plants (Govaerts et al. 2021)) name, *Lophophytum leandri*.

| Organism | Genome length (bp) | AT content (%) | Number of genes | NCBI accession code | Article |
| --- | --- | --- | --- | --- | --- |
| *Arabidopsis thaliana* | 154,478 | 63.7 | 119 | NC_000932.1 | Sato et al. 1999 |
| *Solanum lycopersicum* | 155,461 | 62.1 | 126 | NC_007898.3 | Kahlau et al. 2006 |
| *Populus trichocarpa* | 157,033 | 63.3 | 127 | NC_009143.1 | Tuskan et al. 2006 |
| *Medicago truncatula* | 124,033 | 66.0 | 114 | NC_003119.8 | Unpublished |
| *Erythranthe guttata* | 153,168 | 62.3 | 120 | NC_068035.1 | Zhao et al. 2023 |
| *Oryza sativa* | 134,502 | 61.0 | 129 | NC_031333.1 | Kim et al. 2015 |
| *Phalaenopsis equestris* | 148,959 | 63.3 | 115 | NC_017609.1 | Jheng et al. 2012 |
| *Rhopalocnemis phalloides*, sample MLP | 18,347 | 84.2 | 16 | PQ849603.1 | Unpublished |
| *Rhopalocnemis phalloides*, sample WS | 18,970 | 86.2 | 16 | PQ849602.1 | Unpublished |
| *Rhopalocnemis phalloides*, sample DWS | 19,137 | 86.6 | 16 | MZ269413.1 | Yu et al. 2022 |
| *Rhopalocnemis phalloides*, sample 2194 | 18,174 | 84.3 | 16 | PX369855.1 | This article |
| *Rhopalocnemis phalloides*, sample 3451 | 19,430 | 86.9 | 16 | PX369856.1 | This article |
| *Rhopalocnemis phalloides*, sample 1387 | 18,832 | 86.8 | 16 | MK036331.1 | Schelkunov et al. 2019 |
| *Rhopalocnemis phalloides*, sample 1002 | 19,664 | 84.1 | 16 | PX369854.1 | This article |
| *Scybalium depressum* | 16,513 | 84.5 | 16 | PQ766325.1 | Ceriotti et al. 2025 |
| *Corynaea crassa* | 16,644 | 85.4 | 15 | PQ766322.1 | Ceriotti et al. 2025 |
| *Helosis cayennensis* | 15,715 | 85.9 | 14 | PQ766323.1 | Ceriotti et al. 2025 |
| *Lophophytum mirabile*, sample Aguas Negras | 18,900 | 86.6 | 14 | PQ766326.1 | Ceriotti et al. 2025 |
| *Lophophytum mirabile*, sample Bolivia | 19,095 | 86.5 | 14 | PQ766327.1 | Ceriotti et al. 2025 |
| *Lophophytum mirabile*, sample Calilegua 2 | 18,855 | 86.6 | 14 | PQ766328.1 | Ceriotti et al. 2025 |
| *Lophophytum mirabile*, sample Santa Clara | 18,859 | 86.6 | 14 | PQ766329.1 | Ceriotti et al. 2025 |
| *Lophophytum pyramidale*, sample H. A. Sato 438<sup>a</sup> | 20,892 | 88.4 | 14 | MT834848.1 | Ceriotti et al. 2021 |
| *Lophophytum pyramidale*, sample San Ignacio 2 | 20,902 | 88.4 | 14 | PQ766330.1 | Ceriotti et al. 2025 |
| *Ombrophytum subterraneum* | 17,313 | 86.0 | 15 | MT834847.1 | Ceriotti et al. 2021 |
| *Balanophora fungosa* | 15,130 | 87.4 | 20 | MN414176.1 | Chen et al. 2020 |
| *Balanophora harlandii* | 16,056 | 87.1 | 21 | MN414177.1 | Chen et al. 2020 |
| *Balanophora laxiflora* | 15,505 | 87.8 | 19 | KX784265.1 | Su et al. 2019 |
| *Balanophora reflexa* | 15,507 | 88.4 | 19 | KX784266.2 | Su et al. 2019 |
| *Balanophora yakushimensis* | 14,624 | 87.3 | 18 | NC_062820.1 | Unpublished |
| *Langsdorffia hypogaea* | 17,684 | 86.4 | 16 | PQ766324.1 | Cerioti et al. 2025 |
| *Thonningia sanguinea*, sample T01 | 19,013 | 79.6 | 22 | NC_080980.1 | Kim et al. 2023 |
| *Thonningia sanguinea*, sample T02 | 18,571 | 78.8 | 22 | OQ810030.1 | Kim et al. 2023 |
| *Thonningia sanguinea*, sample T03 | 18,560 | 78.8 | 22 | OQ810031.1 | Kim et al. 2023 |
| *Sarcophyte sanguinea*, sample S01 | 28,384 | 80.9 | 27 | NC_080979.1 | Kim et al. 2023 |
| *Sarcophyte sanguinea*, sample S02 | 28,372 | 80.9 | 27 | OQ810028.1 | Kim et al. 2023 |

### Polymorphism of the plastid genomes of rhopalocnemis

On the basis of 7 samples, the MPPI of the rhopalocnemis plastid genome is 68.7%, the MPMLD is 0.52, and the π is 0.21. As we will show further (see the section “Is the plastid genome of rhopalocnemis the most polymorphic plastid genome among flowering plants?”), the plastid genome of rhopalocnemis possesses record polymorphism among flowering plants with known plastid genomes.

All analyses, the results of which are presented in this article, were conducted using multiple alignments performed by MAFFT. To check for possible MAFFT errors, we also performed analyses of the plastid genomes using alignments made by Muscle and PRANK, but their multiple alignments yielded approximately the same values for MPPI, MPMLD, and π (Supplementary Table 3).

The table with pairwise percent identities of the plastid genomes of rhopalocnemis is shown in Figure 2a.

**Figure 2.**
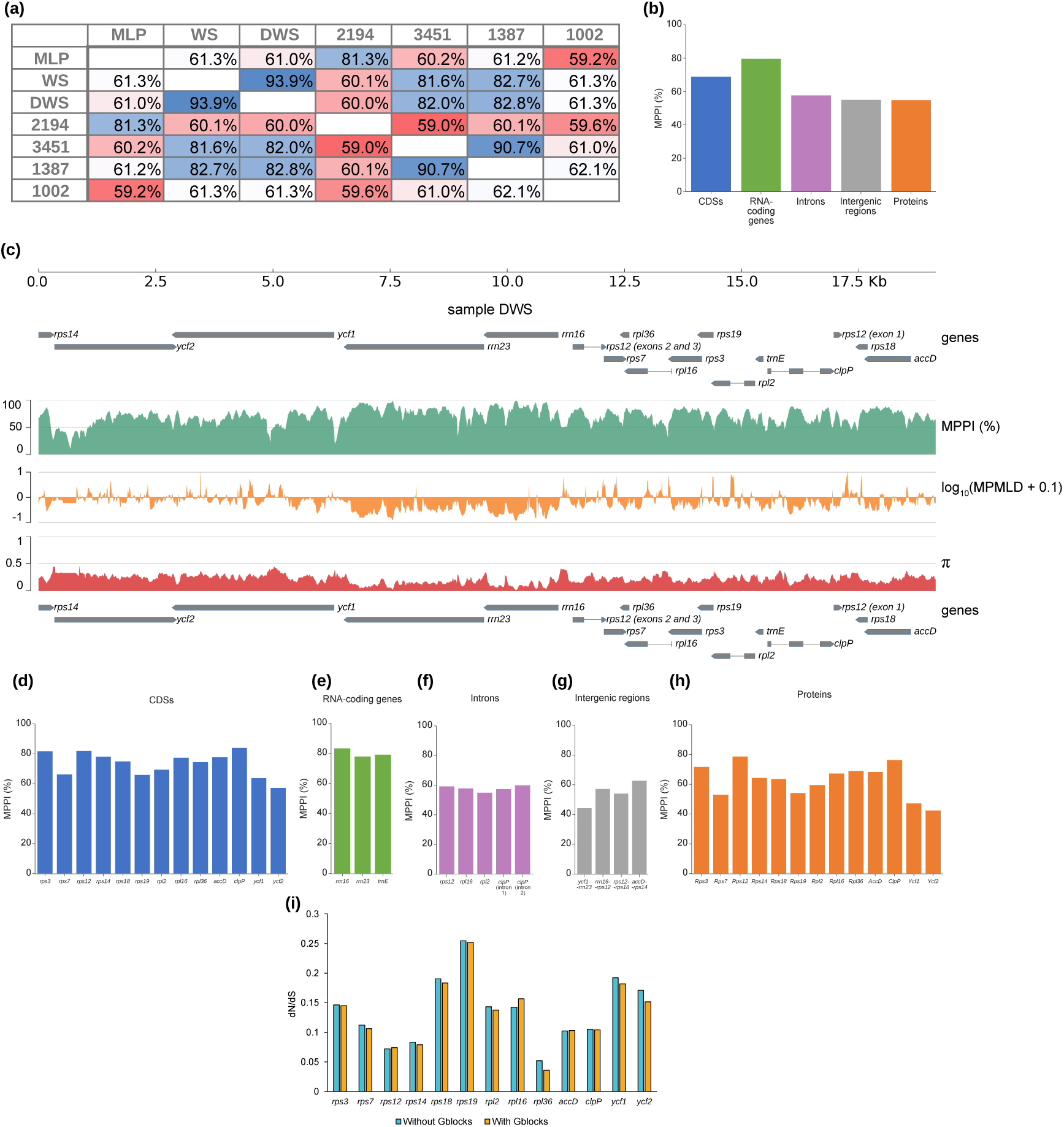
Polymorphism and selection in the plastid genomes of rhopalocnemis. (a) Pairwise percent identities when comparing the plastid genomes of samples of rhopalocnemis entirely. Values are colored from lowest (red) to highest (blue). (b) MPPI in concatenated sequences of different types of regions in the plastid genomes of rhopalocnemis. (c) Plastid genome of rhopalocnemis with the distributions of MPPI, MPMLD, and pi averaged across 100-bp windows. The diagram is made relative to sample DWS but will not differ significantly for the genomes of other samples because of their collinearity and similar length. For convenience, genes are drawn both above and below. When calculating the MPMLD, logarithmic transformation was used to reduce the influence of extreme values; in addition, 0.1 was added to avoid taking the logarithm of 0. (d) MPPI in CDSs. (e) MPPI in RNA-coding genes. (f) MPPI in introns. For introns shorter than 100 bps, the results are not shown because of the low accuracy of calculations for short lengths. (g) MPPI in intergenic regions. For intergenic regions shorter than 100 nucleotides, the results are not shown because of the low accuracy of calculations for short lengths. (h) MPPI in proteins. (i) dN/dS in the CDSs of rhopalocnemis.

Genetic polymorphism is unevenly distributed across the plastid genome of rhopalocnemis. The lowest genetic polymorphism was found in RNA-coding genes, whereas it was somewhat higher in CDSs and even higher in introns and intergenic regions (Figures 2b–2h). For example, the concatenated nucleotide sequences of CDSs from two randomly selected samples of rhopalocnemis are on average only 68.9% similar, while the amino acid sequences of proteins are 54.9% similar.

The genetic polymorphism in the plastid genomes of rhopalocnemis is so high that the plastid genes of two samples often differ from each other more than, for example, the plastid genes of arabidopsis and rice (Figure 1c)—and this despite the fact that rhopalocnemis is a single species, whereas arabidopsis and rice are a eudicot and monocot, respectively.

Despite the high genetic polymorphism, selection on the protein-coding genes of rhopalocnemis is clearly negative (Figure 2i). There was no statistically significant (q-values of t-tests with a Benjamini–Hochberg correction ≤ 0.05) correlation between the intensity of selection and the MPPI, AT content, or CDS length (Figure 3). This may be due not to the absence of correlation but to the small sample size, as rhopalocnemis has only 13 protein-coding genes. The numbers on which Figure 3 was based can be found in Supplementary Table 1.

**Figure 3.**
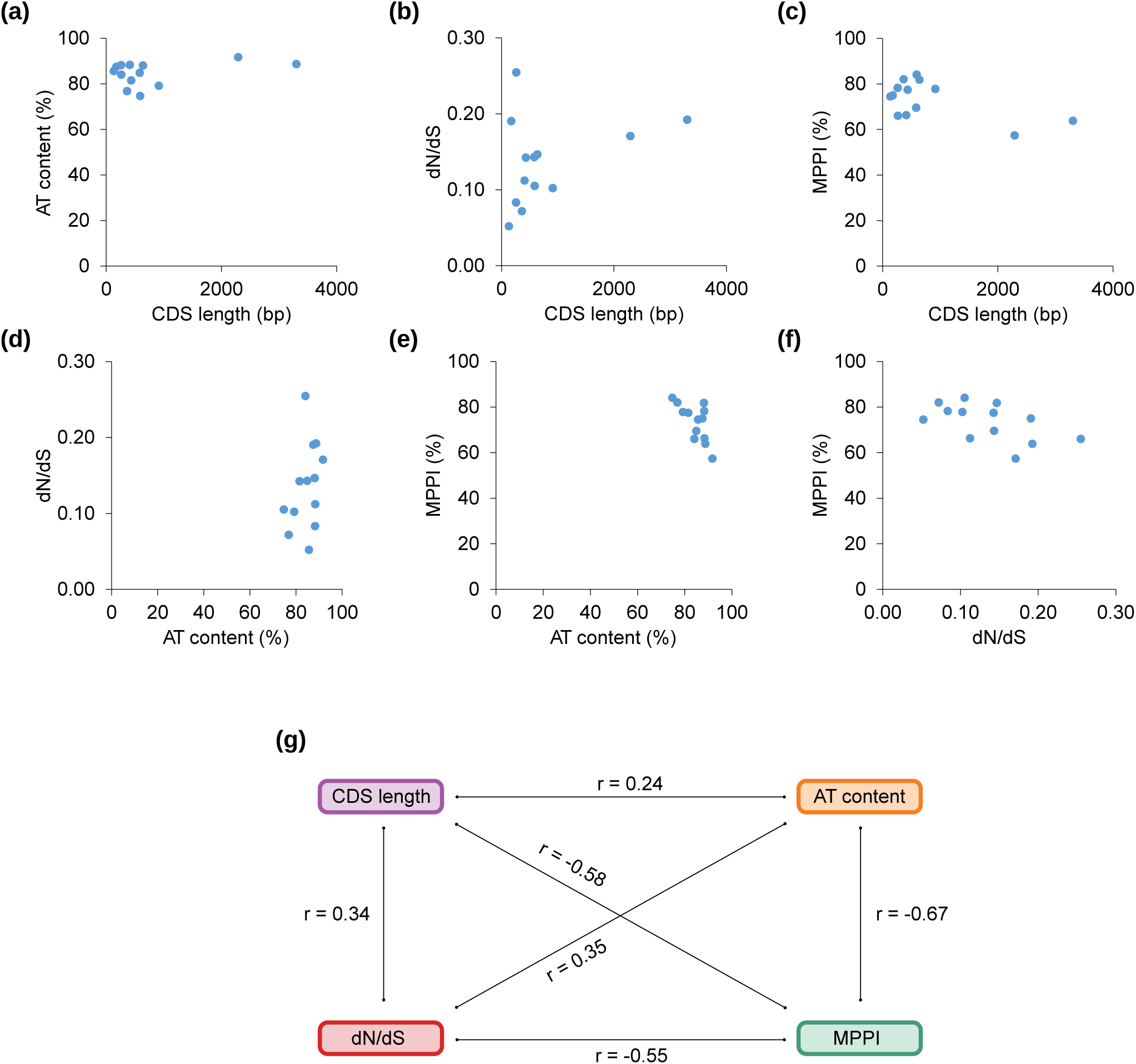
Correlations between the characteristics of the CDSs. (a)–(f) Scatter plots showing dependencies. Each point represents a protein-coding gene. (g) Pearson’s correlation coefficients between characteristics. All correlations are statistically nonsignificant (q-value > 0.05), partly because of the small number of genes.

In the samples that we collected (samples 2194, 3451, 1387, and 1002), we did not observe clear morphological differences that may suggest that these samples belong to different species. Since the samples MLP, WS, DWS cluster on the tree (Figure 1c) with these samples, we do not consider *Rhopalocnemis phalloides* to represent a group of species rather than a single species. Additionally, the fact that plastid genomes on the tree do not group into a small number of clusters, within each of which phylogenetic distances would be low (Figure 1c), supports the idea that this is a single species.

### What happened to *trnE*?

The first article about rhopalocnemis, which studied sample 1387, reported that this species has only one gene encoding tRNA—the leucine tRNA gene *trnL-UAA* (Schelkunov et al. 2019). However, for the other 6 samples of rhopalocnemis, tRNAscan-SE (a standard program for predicting tRNA genes) gave different predictions for the same genomic location:

1. Sample MLP: absence of any tRNA-coding gene.
2. Sample WS: absence of any tRNA-coding gene.
3. Sample DWS: *trnI-UAU*.
4. Sample 2194: absence of any tRNA-coding gene.
5. Sample 3451: *trnL-UAA*.
6. Sample 1387: *trnL-UAA*.
7. Sample 1002: *trnE-UUC*.

It is quite strange for different samples of the same species to have different genes predicted in the same genomic location.

Using multiple alignment of plastid genomes, we identified regions in the samples MLP, WS, and 2194 that corresponded to the tRNA-coding region of the samples DWS, 3451, 1387, and 1002. We compared the sequences of these regions from the 7 samples of rhopalocnemis with all the tRNAs of *Arabidopsis thaliana* and observed that for all 7 samples, the region was most similar to the tRNA gene of the glutamic acid of arabidopsis, namely, *trnE-UUC*. Thus, apparently, all 7 samples of rhopalocnemis have *trnE-UUC*, but tRNAscan-SE recognized this gene differently (and, sometimes, did not recognize it at all) because its sequence is highly mutated.

In photosynthetic plants, *trnE-UUC*, has another function in addition to participating in translation as a tRNA, specifically, it is necessary for the synthesis of tetrapyrroles (Beale 1990). We hypothesized that in rhopalocnemis, *trnE* could have lost its ability to take part in translation and now has only the second function. To assess this possibility, we compared the structures of *trnE* in rhopalocnemis, other Balanophoraceae, and 7 model flowering plants (Figure 4). It is evident that all rhopalocnemis samples have insertions and deletions in the D-arm, and 6 out of 7 samples of rhopalocnemis have mutations that change the anticodon. In addition to rhopalocnemis, anticodon changes were observed in several other species of the Balanophoraceae family.

**Figure 4.**
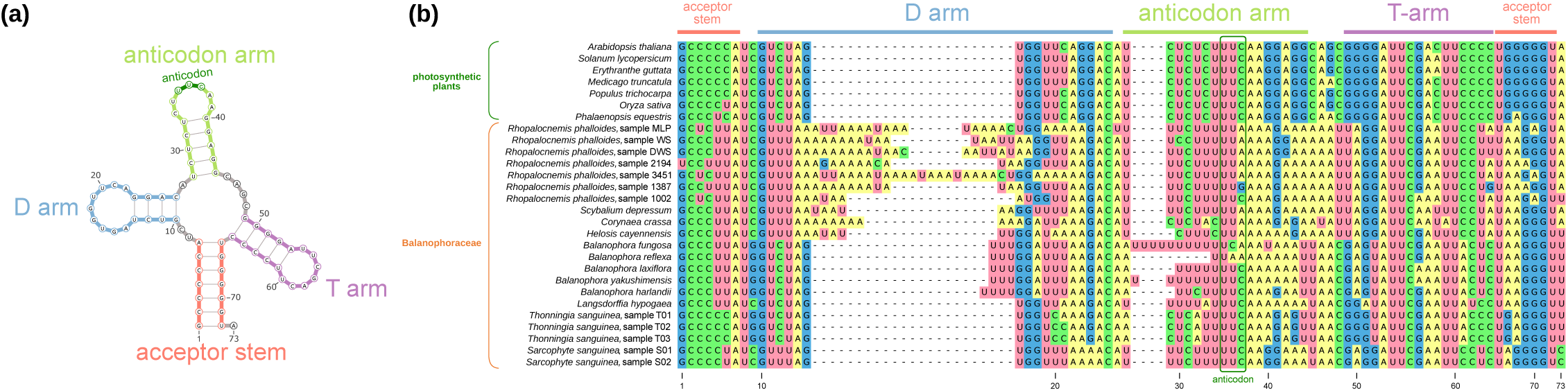
Analysis of *trnE* in Balanophoraceae. (a) Structure of the plastid *trnE* in *Arabidopsis thaliana*. It is the typical cloverleaf structure. (b) Multiple alignment of plastid *trnE* sequences from seven model flowering plants, as well as all Balanophoraceae species in which *trnE* is encoded in plastids. Regions corresponding to different tRNA elements are marked.

At the same time, the nucleotide sequences of the acceptor stem and T-arm in rhopalocnemis are very similar to the corresponding sequences in arabidopsis. The genetic polymorphism in the *trnE* gene of rhopalocnemis is approximately the same as that in other genes and lower than that in intergenic regions and introns (Figure 2), indicating that the *trnE* gene in rhopalocnemis is under negative selection.

It is unknown which specific regions of *trnE* in plants are necessary for functioning in the tetrapyrrole synthesis pathway. However, significant mutations in the D-arm and anticodon, with relative conservation of the acceptor stem and T-arm, suggest that this gene of rhopalocnemis (and several other Balanophoraceae) has ceased to function as a tRNA, but judging by the observed negative selection, it continues to participate in the tetrapyrrole synthesis pathway.

### Is the AT content of the plastid genome of rhopalocnemis still increasing?

One of the unusual properties of plastid genomes of Balanophoraceae is their extremely high AT content, which is approximately 80–90% in most studied species (Table 2). The plastid genomes of Balanophoraceae are among the most AT-rich genomes known to science. It is hypothesized that the increase in AT content arose from the same cause as the increased rate of mutation accumulation and, consequently, the increased genetic polymorphism—namely, the loss of some plastid genome repair genes encoded in the nuclear genome—although the reason for this loss itself is unknown (Schelkunov et al. 2021; Ceriotti et al. 2022; Chen et al. 2023).

One of the questions that interested us was whether the increase in AT content stopped or is still ongoing. To answer this question, in this study, we reconstructed the ancestral sequences of the plastid genomes of rhopalocnemis. To our surprise, the reconstruction results show that in the past, the AT content of the plastid genomes of rhopalocnemis was *even higher* than it is now, and it is currently decreasing (see Supplementary Note 2 for details). To check whether this is an artifact caused, for example, by incorrect reconstruction of ancestral sequences due to long branch lengths, we conducted 1,000 simulations in which a sequence with a similarly high AT content as that in the studied samples evolved on a tree with the same topology as that of the actual phylogenetic tree of rhopalocnemis. In none of these 1,000 simulations did we observe a decrease in the AT content as strong as we see in rhopalocnemis (see Supplementary Note 2 for details). Therefore, it is quite possible that the AT content in rhopalocnemis is indeed decreasing.

It is possible that some time after the plastid genome repair genes were lost (which might have led to harmful mutations), the mechanisms of plastid genome repair were strengthened again, resulting in a reduction in AT content.

Nevertheless, we cannot rule out that the decrease in AT content we observe is simply an artifact of unknown origin.

### Is the plastid genome of rhopalocnemis the most polymorphic plastid genome among flowering plants?

Considering the high polymorphism in the plastid genome of rhopalocnemis, we sought to determine whether it is the most polymorphic among all plastid genomes of flowering plants. For comparison, we used all plastid genomes of species that had at least two plastid genomes on the NCBI website as of 2025-07-09.

During the analysis, we discovered that some plastid genomes in the NCBI databases do not belong to the declared plant. For example, the plastid genome with accession code MZ323108.1 is labeled as belonging to *Arabidopsis thaliana*, but even a simple BLAST analysis indicates that this plastid genome belongs to some plant in the legume family.

Such taxonomically misclassified plastid genomes may cause overestimation of the level of intraspecific polymorphism in the species they are ascribed to; to address this problem, we undertook a search for misclassified plastid genomes in the NCBI databases. The details of the method that we used are described in the Materials and Methods, but in brief, the basis of our method was that we found the *rrn16* and *rrn23* genes in all plastid genomes and constructed a tree based on these genes—if a genome did not fall within the clade of the taxon where it should be, we considered that this genome was likely misclassified. Our method is optimized (see details in Materials and Methods) to effectively identify cases where a plastid genome was mistakenly ascribed to a species that is taxonomically distant from the actual species or to a species whose plastid genome sequences differ significantly from the actual species. In other cases, our method might not detect misclassification—but for finding the most polymorphic plastid genomes, our method is sufficient.

We identified 60 cases in which the plastid genome was misclassified (Supplementary Table 4). Additionally, we found 16 plastid genomes with assembly errors in which the *rrn16* or *rrn23* genes were missing from the plastid genome assembly (Supplementary Table 5); we consider such cases to be assembly errors because these genes are indispensable, and even in non-photosynthetic plants with highly reduced plastid genomes, these genes are always present. Furthermore, we discovered 4 cases in which, owing to an assembly error, the *rrn16* or *rrn23* genes of bacteria, rather than those of plants, were included in the plastid genome assembly (Supplementary Table 6); this issue was previously mentioned in a 2022 article by Robinson et al., but as we found, new plastid genomes with this problem have since appeared on the NCBI website. We reported the 80 aforementioned issues to the authors of these plastid genome assemblies when we were able to find an author’s email. All these problematic plastid genomes were excluded from the analysis of polymorphism.

In addition to the three types of errors mentioned above, we also encountered an error in which the sequence of plastid genomes is a concatenation of several plastid sequences. For example, for *Magnolia patungensis*, there are 3 plastid genomes in the NCBI databases, with lengths of 160,120 base pairs (accession code MZ675531.2), 160,139 base pairs (accession code OP689708.1), and 345,184 base pairs (accession code NC_066227.1). The third plastid genome is incorrectly assembled and is a concatenation of several (not an integral number of) plastid genomes of this species. Errors of this kind lead to an underestimation of the percent identity between plastid genomes; as such, an “extra” sequence will be considered a very long indel. We did not conduct a systematic search for such problems, but to mitigate their effect, we simply processed all multiple alignments of plastid genomes with Gblocks. Gblocks removes dubiously aligned regions, thereby eliminating their effect on polymorphism assessment. Since a highly polymorphic region also may seem dubious to Gblocks, the use of Gblocks may lead to an underestimation of polymorphism (i.e., overestimation of MPPI and underestimation of MPMLD and π). Therefore, the results provided in Supplementary Table 7 cannot be directly compared with the values of polymorphism provided in all other parts of this article. However, since Gblocks affects the results of polymorphism analysis in all plastid genomes in the same way, all the numbers within Supplementary Table 7 can still be compared with each other.

Among all the plastid genomes of flowering plants, the plastid genome of rhopalocnemis was the most polymorphic in terms of MPPI, MPMLD, and π (Supplementary Table 7).

The 10 species with the lowest MPPI (after Gblocks processing), with the MPPI values listed in ascending order, are as follows:

1. *Rhopalocnemis phalloides*: 82.3%.
2. *Monotropa hypopitys*: 94.7%.
3. *Macrosolen cochinchinensis*: 94.8%.
4. *Epipogium roseum*: 95.0%.
5. *Cuscuta gronovii*: 95.4%.
6. *Halerpestes tricuspis*: 95.7%.
7. *Carex littledalei*: 95.7%.
8. *Silene jeniseensis*: 95.7%.
9. *Gentiana leucomelaena*: 95.8%.
10. *Suaeda salsa*: 95.8%.

Of these 10 species, 3 are non-photosynthetic and parasitic: *Rhopalocnemis phalloides, Monotropa hypopitys*, and *Epipogium roseum. Monotropa hypopitys* and *Epipogium roseum* parasitize fungi, unlike *Rhopalocnemis phalloides*, which parasitizes plants. Additionally, 2 species are mixotrophic, meaning that they can both photosynthesize and parasitize: *Macrosolen cochinchinensis* and *Cuscuta gronovii*.

Thus, the 5 most polymorphic plastid genomes belong to species that are heterotrophic or mixotrophic. This finding is related to the previously known high rate of mutation accumulation in heterotrophic and mixotrophic plants (Bromham et al. 2013). The reason for such a high rate of mutation accumulation in heterotrophic and mixotrophic plants is associated with the loss of nuclear genes responsible for the repair of the plastid genome (Schelkunov et al. 2018, 2021; Ceriotti et al. 2022; Chen et al. 2023; Timilsena et al. 2023; Guo et al. 2024). The exact reason natural selection favors this loss is unknown, but several hypotheses have been proposed (Schelkunov et al. 2021).

Notably, out of approximately 300,000 (Christenhusz & Byng 2016) species of flowering plants, only for approximately 5,000 species is the plastid genome sequence known from more than one sample (Supplementary Table 7), thus allowing an evaluation of polymorphism. It is quite possible that there are species of flowering plants with a higher polymorphism of the plastid genome than that of rhopalocnemis. It is possible that such species exist even within the Balanophoraceae family, as the increase in the rate of mutation accumulation is characteristic, to some extent, of all Balanophoraceae with known plastid genome sequences (Figure 1c).

On the other hand, it should be noted that in nature, rhopalocnemis has been observed from Nepal to the Moluccas (a distance of approximately 6,000 kilometers; Govaerts et al. (2021)), while the maximum distance between samples studied in this study is approximately 1,300 kilometers (Figure 1b). Therefore, the polymorphism of rhopalocnemis in our study may even be *underestimated*.

### Is the plastid genome of rhopalocnemis the most polymorphic among all genomes?

As of 2025, the most polymorphic of all genomes of all taxa is considered the nuclear genome of the basidiomycete fungus *Schizophyllum commune* (Baranova et al. 2015). The MPPI in the whole-genome analysis of the nuclear genomes of 13 samples of *Schizophyllum commune* is 83.6% (Marian et al. 2024). In the whole-genome analysis of the plastid genomes of rhopalocnemis, the MPPI is 68.7%.

Notably, *Schizophyllum commune* reproduces sexually, so its high polymorphism can lead to the following problem: if two proteins work interacting with each other (for example, they are parts of a protein complex), and one protein is encoded in the maternal haplotype and the other in the paternal haplotype, then owing to high polymorphism, these proteins may be poorly compatible with each other in the sense that their contact will be unstable.

In plants (including, probably, rhopalocnemis), plastid genomes are usually inherited from only one parent (Schneider 2023), so there is no problem of incompatibility between plastid proteins. However, the products of all 16 genes encoded in the plastid genome of rhopalocnemis work in contact with the products of genes encoded in the nuclear genome. For example, 11 out of 16 genes in the plastid genome of rhopalocnemis encode components of the plastid ribosome, but at least 30 other components (Schelkunov et al. 2021) of the plastid ribosome are encoded in the nuclear genome of rhopalocnemis. Thus, the high polymorphism of the plastid genome can complicate the joint functioning of proteins or RNAs encoded in different organelles.

The polymorphism in the nuclear genome of rhopalocnemis is currently unknown. The rate of mutation accumulation in the nuclear genome, as determined by the study of the transcriptome, is several times greater than that in photosynthetic plants (Schelkunov et al. 2021)—but this difference is much lower than that in its plastid genomes (Figure 1c); thus, it is likely that the nuclear genome of rhopalocnemis is not as polymorphic as the plastid genome.

## Conclusions

In this study, we demonstrated that the high mutation accumulation rate in the plastid genome of rhopalocnemis, associated with the loss of nuclear genes responsible for plastid genome repair, led to a very high level of polymorphism. At the same time, selection on the genes encoded in the plastid genome is negative. The peculiarity of the plastid genome of rhopalocnemis also lies in the alteration of the tRNA of glutamic acid, which has ceased to partake in translation and now performs a different function.

Additionally, while conducting this study, we discovered several dozen cases of considerable errors in the plastid genomes deposited in the NCBI databases. Accounting for these errors may improve the results of other scientific studies.

## Materials and Methods

### Sample collection, sample preparation, and sequencing

Samples 2194, 3451, and 1002 were collected in Vietnam during the Vietnam–Russia Tropical Center expeditions. Information about the collection sites is provided in Supplementary Table 8.

DNA was extracted from silica gel-dried or ethanol-fixed samples using a modified CTAB method (the modifications were as follows: polyvinylpyrrolidone was added to the CTAB buffer, and chloroform extraction was performed twice) (Doyle 1987). For the ethanol-fixed material, we first rinsed the samples several times in distilled water and then dried them. DNA was sheared using a Covaris S220 instrument and then purified using 0.9x volume of AMPure magnetic beads. Libraries were prepared using a NEBNext Ultra II library preparation kit. Sample 3451 was sequenced using Illumina NovaSeq 6000 instrument, while samples 2194 and 1002 were sequenced using an Illumina NextSeq 500 instrument. For sample 3451, paired-end reads of 151 bps were obtained, and for samples 2194 and 1002, paired-end reads of 76 bps were obtained.

The numbers of produced reads (a pair of reads is counted as two reads) were as follows:

1. Sample 2194: 364,591,350.
2. Sample 3451: 1,191,575,944.
3. Sample 1002: 596,975,726.

### Plastid genome assembly

We assembled plastid genomes not only for samples 2194, 3451, and 1002 sequenced in this work but also for samples MLP, WS, DWS, and 1387, for which assemblies were available in NCBI before our work. This was done to control the correctness of the assemblies of samples MLP, WS, DWS, and 1387. Illumina read sets of samples MLP, WS, DWS, and 1387 were taken from the SRA database and have identifiers SRR31637715, SRR31637718, SRR14800609, and SRR7995544, respectively.

Before assembly, read trimming was performed using Fastp 0.23.4 (Chen et al. 2018) by sequentially executing the following operations:

1. All bases from the 3’-end of the reads with a Phred score less than 3 were removed.
2. If there were 5 consecutive bases in a read with an average Phred score less than 15, those bases and the entire part of the read toward the 3’-end were removed.
3. Reads in which, after the previous operations, the average Phred score was less than 20 were removed.
4. Reads whose length became less than 30 bases after the previous operations were removed.
5. Adapters were removed by palindromic trimming without specifying the adapter sequences.

The popular assembly programs GetOrganelle (Jin et al. 2020) and SPAdes (Bankevich et al. 2012) were unable to assemble the plastid genomes of all the samples into circular contigs; thus, we created our own assembly pipeline, which works as follows:

1. Assembly is performed using Minia 3 (Chikhi & Rizk 2013). The advantage of Minia is that although it makes quite fragmented assemblies, it works very fast.
2. Alignment of BLASTN 2.11.0+ (Camacho et al. 2009) sequences of closely related plastid genomes to contigs assembled in step 1 is performed. The e-value is allowed to be no greater than 0.001. This step is necessary to approximately determine the contigs related to the plastid genome. As “closely related plastid genomes”, we used the plastid genomes of samples DWS and 1387 because when we started this study, the plastid genomes of samples MLP and WS were not yet available on the NCBI website.
3. The pipeline takes those contigs that had statistically significant matches to the reference in step 2. Then, a script aligns reads to them using BWA-MEM 0.7.17 (Li & Durbin 2009).
4. For those pairs of reads where only one read from the pair aligned, the pipeline takes the second read from the pair.
5. The pipeline uses Unicycler 0.4.9 (Wick et al. 2017) to assemble the reads from step 4. Unicycler is a higher-quality but slower assembler than Minia. Since there are few reads at this stage, speed is not highly important. One of the advantages of Unicycler compared with many assemblers is that if a circular sequence is obtained, it correctly “circularizes” it, meaning that it outputs circular sequences without extra base pairs at the beginning and end—after the last base pair, Unicycler places the first base pair.
6. The pipeline repeats steps 2–5 until the length of the longest plastid contig has stopped changing.

If describing the pipeline briefly, it first performs a rough assembly and then iteratively incorporates reads such that at least one read from a pair aligns to the assembly, creating a new assembly at each iteration. Overall, the scheme of this pipeline is similar to that of the GetOrganelle pipeline, but this pipeline is better suited for assembling plastid genomes with a high mutation accumulation rate. The reason for this is that GetOrganelle initially attracts reads for assembly that align to the reference using Bowtie2 (Langmead & Salzberg 2012), whereas this pipeline attracts Minia contigs that align to the reference using BLAST; the advantage of BLAST for this task is that it aligns dissimilar sequences better than Bowtie2 does.

The correctness of the assemblies was verified using alignment of reads to the assembled plastid genomes as described below. Since the coverage by reads at the very end and the very beginning of a plastid genome decreases because of difficulties in aligning reads that cross the edges of a circular contig, all the operations listed below were performed not only for the original version of the plastid contig but also for a version of the plastid contig in which the first 10,000 bps were moved to the end, thereby changing the closure point of the circle. To verify the correctness of the plastid genome assemblies, we performed the following operations:

1. We aligned the reads to the plastid genome using BWA-MEM 0.7.17.
2. Sequences similar to the plastid genome may be found in the nuclear and mitochondrial genomes. To minimize the alignment of mitochondrial and nuclear reads to the plastid genome, we used custom scripts to retain only reads that had at least 50 bps aligned, and in which the similarity to the plastid genome in the aligned part was at least 98%.
3. Since the AT content affects the coverage by Illumina reads, we adjusted the coverage for AT content using the correctGCBias program from the DeepTools 3.5.5 suite (Ramírez et al. 2016).
4. We visualized the coverage distribution across the alignment made in step 3 using the Plotly library (Plotly Technologies, Inc 2015) of the Python programming language to ensure that there were no regions with abnormally high or low coverage. Additionally, to verify the proper functioning of correctGCBias, we visualized the AT content distribution and the coverage distribution before the correction was made by correctGCBias. All the graphs can be found in Supplementary Note 1.
5. We calculated the minimum and maximum coverage of the plastid genome both before and after correction to programmatically verify the accuracy of the visual analysis results.

To compare the new assemblies of plastid genomes of samples MLP, WS, DWS, and 1387 with the assemblies of these samples available on the NCBI website, we performed pairwise alignments using the online version of BLASTN. The BLAST results differed for the WS and 1387 samples. Coverage analysis demonstrated that the assemblies made in this study are correct, rather than those on the NCBI website; see Supplementary Note 1 for details. The sequence of sample 1387 was previously uploaded to the NCBI website by us, and now we replaced the sequence with a more accurate one. We also notified the authors of the plastid genome assembly of the sample WS that it contains an error.

For samples MLP, WS, and DWS, the SRA database contains not only Illumina reads but also PacBio reads. For samples MLP and WS, these are high-accuracy PacBio HiFi reads, and for sample DWS, these are low-accuracy PacBio CLR reads. We verified the assemblies through coverage analysis in the same way as for the Illumina reads, but with the following differences:

1. The alignment was performed not with BWA-MEM 0.7.17 but with Minimap 2.28 (Li 2018). For the PacBio HiFi reads, it was done with the --map-hifi option, and for the PacBio CLR reads with the --map-pb option.
2. For PacBio HiFi reads, we required at least 10,000 bases to align with a minimum similarity of 98%. For PacBio CLR reads, we required at least 10,000 bases to align with a minimum similarity of 90%. As with the Illumina reads, filtering by the length of the aligned part and minimum similarity was performed using our own scripts.

The alignment of PacBio reads confirms the accuracy of our assembly variant of the plastid genome of sample WS (see Supplementary Note 1).

### Annotation of the plastid genomes

Annotation was performed not only for the plastid genomes of the three samples sequenced for this work but also for the other four samples to verify the accuracy of the annotations posted on the NCBI website.

The annotation included the following steps:

1. TBLASTN 2.11.0+ alignment of reference proteins to the plastid genomes was performed with allowed e-values no higher than 10^-3^. The reference included all proteins of the *Arabidopsis thaliana* plastid genome (taken from the NCBI annotation with the accession code NC_000932.1), plus the *InfA* protein from the *Santalum album* plastid genome (taken from the NCBI annotation with the accession code MN106256.1), as well as proteins from the plastid genomes of the samples DWS and 1387 of rhopalocnemis (the only samples whose annotations were present on the NCBI website at the start of our work). *InfA* was taken from *Santalum album* because it is the only plastid-encoded protein whose gene is present in the plastid genomes of almost all flowering plants but is absent in *Arabidopsis thaliana. Santalum album* was used since it is a close relative of rhopalocnemis that has not lost this gene. *Arabidopsis thaliana* was used because it is a model plant, which reduces the likelihood of annotation errors in its genome.
2. BLASTN 2.11.0+ alignment of reference rRNAs and tRNAs to plastid genomes was performed with allowed e-values no higher than 10^-3^. Reference rRNAs and tRNAs were taken from *Arabidopsis thaliana*, as well as from samples DWS and 1387.
3. Using the getorf program from the EMBOSS 6.5.7 software suite (Rice et al. 2000), all open reading frames that were at least 60 bps in length were identified. As an open reading frame, we considered a region between two stop codons that does not contain stop codons.
4. For the open reading frames identified in step 3, similarity with models from the Pfam-A database (Mistry et al. 2021) version 35.0 was searched for using the hmmscan program from the HMMER 3.2.1 software suite (Mistry et al. 2013). E-values no higher than 10^-3^ was allowed.
5. Alignment of the open reading frames identified in step 3 was performed using BLASTP 2.11.0+ to proteins of the NCBI nr database (https://ftp.ncbi.nlm.nih.gov/blast/db/) as of 2023-04-09. E-values no higher than 10^-3^ were allowed.
6. A search for RNA-coding genes (i.e., those coding for rRNA, tRNA, and other RNAs that do not encode proteins) was conducted by scanning the plastid genomes with Infernal 1.1.5 (Nawrocki & Eddy 2013) using models from the Rfam 14.9 database (Kalvari et al. 2021). E-values no higher than 10^-3^ were allowed.
7. A search for tRNA-coding genes was conducted using tRNAscan-SE 2.0.12 (Chan & Lowe 2019) with the “organelle” tRNA model (the option “-O”).

The results of the methods described above were analyzed manually.

The methods described above allowed the discovery of previously unnoticed *trnE* genes in samples MLP and WS. We informed the authors of the annotations of those two samples about this.

Additionally, for the analysis of the *trnE* gene (see the section “What happened to *trnE*?”), we used the following methods:

1. To ensure that the tRNA gene we observed in rhopalocnemis originated from the *trnE* gene, we first performed a multiple alignment of this gene from all seven samples of rhopalocnemis using MAFFT 7.520 (Katoh & Standley 2013) with the options “--maxiterate 1000 –globalpair”. Then, on the basis of the obtained alignment, we created a Markov model using the hmmbuild program from the HMMER 3.3.2 software suite. Afterward, we analyzed the similarity of all *Arabidopsis thaliana* tRNAs with this model using the nhmmer program from the HMMER 3.3.2 software suite to determine which tRNA of *Arabidopsis thaliana* the tRNA gene of rhopalocnemis most closely resembles.
2. To identify *trnE* that may have been unnoticed because of its nonstandard sequence, in all the plants of the Balanophoraceae family except rhopalocnemis, we scanned all the plastid genomes of Balanophoraceae presented in Table 2 using the Markov model from step 1. This method allowed us to find a previously unnoticed *trnE* in the plastid genome of *Corynaea crassa*. We reported this to the authors of the annotation of that genome.
3. To find bases homologous between *trnE* of Balanophoraceae and of model photosynthetic plants (Figure 4b), we performed a multiple alignment of *trnE* of these species using MAFFT with the options “--maxiterate 1000 --globalpair”.

To refine the annotation of intron boundaries, we used RNA-seq reads. The RNA-seq reads were available in the SRA for samples MLP, WS, DWS, and 1387. The SRA identifiers were as follows:

1. Sample MLP: SRR31637714.
2. Sample WS: SRR31637717.
3. Sample DWS: SRR14800310.
4. Sample 1387: SRR7995545, SRR12771436, SRR12771435. We concatenated the reads from these three sets.

The RNA-seq reads were trimmed using Fastp in the same way as the genomic reads were (see the beginning of the section “Plastid genome assembly” for the description of the trimming method). The RNA-seq reads were aligned to the corresponding plastid genomes using STAR 2.7.3a (Dobin et al. 2013) with the option “--genomeSAindexNbases 6”. Subsequently, we visualized the RNA-seq read alignment in CLC Genomics Workbench 22.0.1 (QIAGEN), which allowed us to determine the intron boundaries. The annotation of intron boundaries in the remaining three samples (samples 2194, 3451, and 1002) was performed considering the annotation of intron boundaries in samples MLP, WS, DWS, and 1387. To do this, we performed a multiple alignment of the plastid genomes of the 7 samples by using MAFFT with the options “--maxiterate 1000– globalpair” and visualized the multiple alignment in CLC Genomics Workbench.

### Methods of polymorphism calculation

A description of the polymorphism metrics is given in Table 1. To calculate the polymorphism metrics, we created a script available at https://github.com/shelkmike/Polymorphism_suite. It takes a multiple alignment as input and outputs the following information:

1. Three tables, each containing all values of percent identity, phylogenetic distance, and p-distance for all possible pairwise comparisons.
2. The values of three polymorphism metrics —MPPI, MPMLD, and π—are arithmetic means of all the values from the tables in step 1.

When the percent identity and p-distance for a pair of sequences in a multiple alignment are calculated, if both sequences have a gap in a given column, that column is completely excluded from the comparison of these two sequences.

It may happen that in a multiple alignment, two sequences do not overlap at all, meaning that for all columns where one sequence has a base, the other has a gap, and for all columns where the second sequence has a base, the first has a gap. In such cases, the calculation of the percent identity and p-distance is complicated. Therefore, in such cases, the script records the percent identity as 0% and the p-distance as 1.

To calculate phylogenetic distances, the script uses IQ-TREE 1.6.12 (Minh et al. 2020) with default parameters.

The script can perform analysis not only when given multiple alignments as input but also when given pairwise alignments. Since IQ-TREE is not capable of creating a phylogenetic tree from pairwise alignments, our script converts the pairwise alignment into an artificial multiple alignment by simply duplicating the second sequence in the input file. It then creates an IQ-TREE tree and subsequently removes one of the copies of the second sequence from the resulting tree.

The script can perform analyses for both nucleotide sequence alignments and amino acid sequence alignments.

CDS alignments of rhopalocnemis for the polymorphism analysis were made using TranslatorX 1.1 (Abascal et al. 2010). TranslatorX is a program that aligns protein-coding sequences by first aligning the corresponding protein sequences. We ran TranslatorX so that it used MAFFT for alignment. In doing so, we modified the TranslatorX code to run MAFFT with the options “--maxiterate 1000 --globalpair”, which increases alignment accuracy.

Alignments of all other nucleotide sequences, as well as amino acid sequences, were made directly using MAFFT 7.520 with the options “--maxiterate 1000 --globalpair”.

Additionally, to test the possible influence of the multiple alignment tool on the results of the polymorphism analysis, in addition to using MAFFT, we performed alignments of entire plastid genomes using Muscle 5.3 (Edgar 2004) with default parameters, as well as the PRANK (Löytynoja 2014) version posted on GitHub on 2017-04-27, with default parameters. The polymorphism metrics calculated from alignments made with Muscle and PRANK were approximately the same as those from the alignment made with MAFFT (Supplementary Table 3). Therefore, Muscle and PRANK were not used in further analyses.

Except where explicitly stated, the polymorphism analyses for the sequences of rhopalocnemis were conducted on alignments not processed by Gblocks (Castresana 2000). This is because a visual analysis of the multiple alignment did not reveal any clearly misaligned sites, and using Gblocks could lead to the erroneous removal of correctly aligned highly polymorphic sites.

When calculating polymorphism in introns and intergenic regions, we did not use sequences shorter than 100 bases. This is because the calculation of polymorphism for very short sequences is inaccurate. For example, the intergenic region between the *rpl16* and *rps3* genes in the sample DWS is 1 bp long. There are no CDSs or RNA-coding genes shorter than 100 bps in rhopalocnemis, except for the *trnE* gene, whose lengths range from 77 to 96 bps in different samples. We used the *trnE* gene in the analysis of RNA-coding genes’ polymorphism.

### Phylogenetic and natural selection analyses

To construct the tree presented in Figure 1c, the following operations were performed sequentially:

1. For all the species listed in Table 1, the CDSs and RNA-coding gene sequences were taken from the annotations.
2. The alignment of each gene’s CDS was performed separately using TranslatorX, which ran MAFFT with the options “--maxiterate 1000 --globalpair”.
3. The alignment of each RNA-coding gene was performed separately using MAFFT with the options “--maxiterate 1000 --globalpair”.
4. From the alignments made in steps 2 and 3, dubious sites were removed using Gblocks 0.91b. For CDSs, Gblocks was run with the options “-t=c -b3=4”, and for RNA-coding genes, the options “-t=d -b3=4” were used. The “-t” option indicates whether the nucleotide sequence is protein-coding or not, and “-b3=4” increases the stringency in removing dubious sites. The option “-b3=4” was chosen because, with this option, the local version of Gblocks works the same as the online version of Gblocks when the online version is run with its most stringent parameters.
5. The alignments of all the CDS and RNA-coding genes were concatenated using geneStitcher (https://github.com/ballesterus/Utensils), the latest version available on GitHub as of 2025-01-21.
6. The tree was constructed using IQ-TREE 1.6.12, utilizing 1000 pseudoreplicates generated by the UFBoot method, and automatic substitution model selection using the “MFP+MERGE” method. For the model selection, a partitioning of the alignment into CDS and RNA-coding genes was provided to IQ-TREE. The topology at the family level was fixed and made identical to the topology from the article of Li et al. (2021). That is, IQ-TREE determined the topology only within Balanophoraceae.

Analysis of natural selection in the CDSs of rhopalocnemis was conducted using the Codeml program from the PAML 4.10.7 package (Yang 2007) on the basis of alignments made by TranslatorX. Codeml was run with the following parameters:

1. Columns in which at least one sample had a gap were removed.
2. The F3X4 codon model was used.
3. The topology of the rhopalocnemis tree was extracted from the tree that was constructed as described above.
4. The ratio of the transition frequency to the transversion frequency was determined automatically by Codeml. A starting value of 2.0 was used.
5. The dN/dS ratio was determined automatically by Codeml. A starting value of 0.5 was used.
6. Since the goal was to calculate the average dN/dS across samples, Codeml was instructed that each gene has the same (gene-specific) dN/dS across the entire rhopalocnemis tree, and this dN/dS is the same for all codons. In Codeml terms, this model is called M0.

### Analysis of polymorphism in plastid genomes of all flowering plants available on the NCBI website

First, we downloaded all the plastid genomes of flowering plants from the NCBI website available as of 2025-07-09.

For this step, the esearch program from the Entrez Direct software suite version 23.6 (Schuler et al. 1996) was used. The download was performed using the query ‘(“chloroplast” [title] OR “plastid” [title]) AND “flowering plants” [organism] AND 10000:1000000000 [slen]’. The filter for a minimum length of 10,000 bps was used for preliminary (more precise filtering would follow) filtering, allowing the exclusion of cases in which only a specific region of the plastid genome was sequenced and not the entire genome; such a filter helps reduce download time. Then, from the downloaded plastid genomes, only those that simultaneously met the following conditions were retained:

1. The annotation indicates that the sequence is circular. This is necessary to eliminate cases in which only a specific region of the plastid genome was sequenced but not the entire genome.
2. The species name does not contain “x”. These are hybrids.
3. The species name does not contain “cf.”, which indicates that the authors are not sure about the species identification.
4. The species name does not contain “sp.”, which indicates an unknown species.
5. The species name does not contain “aff.”, which indicates an unknown species that is closely related to another species.

Since some plastid genomes on the NCBI site are available in multiple copies (for example, one in the GenBank database and another in the RefSeq database), we retained only one copy for all plastid genomes with identical sequences using the rmdup command of SeqKit 2.7.0 (Shen et al. 2024). After deduplication, 36,257 plastid genomes remained.

We subsequently searched for plastid genomes that were presumably misclassified. Excluding such plastid genomes is important for the analysis of intraspecific polymorphism because the assessment of intraspecific polymorphism in a species to which a plastid genome of another, distantly related, species is mistakenly assigned will be overestimated. The search for plastid genomes that were taxonomically misclassified was conducted as follows:

1. Using Infernal and bacterial 16S and 23S models from the Rfam database, we identified 16S rRNA genes (also known as *rrn16*) and 23S rRNA genes (also known as *rrn23*) in all plastid genomes. During the search, we set the maximum e-value cutoff to 10^-3^. Only 16S and 23S genes (but not other plastid genes) were used to make the calculations less time-consuming.
2. If only one of these two genes or neither of them were found in the plastid genome, we considered the plastid genome to be incomplete and excluded it from further analyses. The absence of these genes indicates an incomplete assembly because even in heterotrophic plants with the most reduced plastid genomes, 16S and 23S genes are present in all known plastid genomes, indicating the extreme importance of these genes.
3. It was previously shown that some plastid genomes uploaded to NCBI were assembled incorrectly and contain fragments of bacterial genomes. To identify and exclude such cases, all sequences found in step 1 were aligned to the 16S rRNA and 23S rRNA databases and clustered by 99% similarity. These databases are parts of the SILVA 138.2 database (Quast et al. 2012). The alignment was performed using Megablast 2.11.0+ with a maximum e-value threshold of 10^-5^. If more than half of the top 5 matches in the database were not to plastid genomes, we considered the gene in question to be the result of an assembly error, where the plastid contig is a chimera of plastid sequence and contamination sequence. Plastid genomes in which at least one of the two genes resulted from such an assembly error were excluded from further analyses.
4. Multiple alignment of all 16S sequences was performed using MAFFT with default parameters. Multiple alignment of all 23S sequences was performed in the same way.
5. The multiple alignments of 16S and 23S were concatenated using Phyutility 2.7.1 (Smith & Dunn 2008).
6. A phylogenetic tree was constructed from the concatenated sequences of these genes using IQ-TREE 2.4.0 with the GTR+Gamma model. To reduce computation time, automatic model selection was not performed. The resulting tree in the Newick format is provided in Supplementary File 1.
7. To determine whether a plastid genome is misclassified, we examined the plastid genomes that were closest to it on the tree. If there was a sister clade for a given plastid genome that contained more than one plastid genome, we considered all these sister plastid genomes. If the sister clade contained only one plastid genome, we also considered the clade that was sister to the clade “given plastid genome + sister clade”. Thus, we identified at least two plastid genomes on the tree that are the closest relatives to the given plastid genome. We called such genomes “plastid genomes of comparison”. We then considered the given plastid genome to be misclassified taxonomically if both of the following conditions were simultaneously met: a) All plastid genomes in the comparison belonged to a single order according to the NCBI Taxonomy database (Schoch et al. 2020), current as of 2025-05-07. That is, there are no contradictions within the plastid genomes of comparison that could indicate a classification error in one of them. b) The given plastid genome did not belong to the same order as the plastid genomes of comparison, according to the NCBI Taxonomy database, current as of 2025-05-07.

The analysis was conducted at the order level, rather than at the levels of lower taxonomic ranks (such as families), because the systematics at the family level in some orders is currently unstable and subject to substantial changes. Attempts to analyze at the family level sometimes led to false predictions of misclassifications. For example, the plastid genome of *Mimulus ringens* (NCBI accession code NC_068043.1) clusters with the plastid genomes of the genus *Mimulicalyx* in the tree. According to NCBI Taxonomy, *Mimulus* belongs to the family Scrophulariaceae and *Mimulicalyx* to the family Phrymaceae, whereas according to the Plants of the World Online database (Govaerts et al. 2021), they both belong to the family Phrymaceae. Thus, in this case, the issue is not an error by the authors who attributed the plastid genome to the wrong species but an error in the systematics of the NCBI Taxonomy database.

After removing plastid genomes that contained assembly errors or were assigned to incorrect orders on the NCBI site, we conducted an analysis of intraspecific polymorphism for all species for which at least two plastid genomes remained. MAFFT alignment of entire plastid genomes is not suitable for polymorphism analysis for two reasons:

1. Most plants have a small single-copy region in their plastid genomes, the orientation of which is difficult to determine precisely because it is located between two long copies of an inverted repeat, complicating assembly. Additionally, recombination sometimes occurs between these copies of the inverted repeat, causing the small single-copy region to change direction. Therefore, plastid genomes of the same species, available on the NCBI website, are often noncollinear with each other, which prevents their alignment by programs such as MAFFT that assume collinearity of the sequences being aligned.
2. Some plastid genomes available on the NCBI website have structural assembly errors. For example, for *Magnolia patungensis*, there are 3 plastid genomes on the NCBI website, with lengths of 160,120 base pairs (accession code MZ675531.2), 160,139 base pairs (accession code OP689708.1), and 345,184 base pairs (accession code NC_066227.1). The third plastid genome is incorrectly assembled and is a concatenation of several (and not an integral number of) plastid genomes of this species.

Thus, to study intraspecific polymorphism, it is necessary to use an alignment method that accounts for the possible presence of structural polymorphism or assembly errors. We used the program for multiple genome alignment progressiveCactus 2.9.7 (Armstrong et al. 2020), which considers structural differences. As input, progressiveCactus requires not only sequences but also a phylogenetic tree of species with branch lengths. We constructed a phylogenetic tree of entire plastid genomes using a method that does not require multiple alignment—specifically, using the program Mashtree 1.4.6 (Katz et al. 2019), which involves constructing a tree using the neighbor joining method based on the number of k-mers shared between plastid genomes. We used a k-mer length of 11 bps for Mashtree. In the analysis, we used all k-mers, not just those that appeared in the genome at least 5 times, as is the default for Mashtree.

Subsequently, to remove the multiple alignment created by progressiveCactus (see step 2 above) regions with possible assembly errors, we processed the alignments with Gblocks using the options “-t=d -b3=4”.

Thus, for each species for which at least two plastid genomes were available on the NCBI website, we obtained a multiple (or pairwise, in the case of two genomes) alignment. After this, we calculated the values of MPPI, MPMLD, and π for each species using our script designed for polymorphism analysis (see its description in the “Methods of polymorphism calculation” section).

Since at the time of this analysis, the sequences of the plastid genomes of samples 2194, 3451, and 1002 of rhopalocnemis had not yet been uploaded to the NCBI website (sequences were available only for MLP, WS, DWS, and 1387), we repeated exactly the same analysis for all 7 samples of rhopalocnemis as we did for the plastid genomes from the NCBI website, including the use of Mashtree, progressiveCactus, and Gblocks. It is important to note that since Gblocks may remove not only erroneous alignment sites but also highly polymorphic sites (which are difficult to distinguish from erroneous sites), the use of Gblocks causes underestimation of the values of polymorphism. However, since Gblocks was applied to all plastid genomes (rhopalocnemis as well as all plastid genomes from NCBI), despite the use of Gblocks, the results of this analysis can be used to rank plastid genomes by polymorphism.

As mentioned above, our method for detecting taxonomic misclassifications may not find misclassifications in some cases—for example, if the classification error was between different taxa within the same order. Therefore, after calculating the polymorphism of the plastid genomes for all the species, we conducted an additional analysis aimed at ensuring that the list of the 10 most polymorphic species (those with the lowest MPPI) did not contain species whose plastid genomes were misclassified. For this analysis, we did the following:

1. Without the use of scripts (manually), we examined the taxonomic affiliation of plastid genomes closely related to the one under consideration on the tree presented in Supplementary File 1.
2. Additionally, without using scripts, we looked at the taxonomic affiliation of the most similar sequences according to the alignment of the plastid genome under consideration using Megablast online to the core_nt database.

Using these two methods, we analyzed the taxonomic affiliation of plastid genomes, ensuring that among the 10 species with the lowest MPPI, not a single one had a polymorphism analysis that included even one taxonomically misclassified plastid genome. For all species for which at least 2 correctly classified plastid genomes remained after the removal of incorrectly classified plastid genomes, we redid the polymorphism analysis on the remaining genomes.

### Analysis of AT content evolution

To answer the question “Is the AT content of the plastid genome of rhopalocnemis still increasing?”, the following actions were taken:

1. The plastid genomes of rhopalocnemis were aligned end-to-end using MAFFT.
2. The branch lengths were determined using IQ-TREE based on the alignment obtained in step 1. In this case, the tree topology was fixed to the one obtained as described in the section “Phylogenetic and natural selection analyses”. Thus, IQ-TREE was used only to determine the branch lengths, which are needed for the FastML (Moshe & Pupko 2019) program (see step “3”.
3. The ancestral sequences of rhopalocnemis were reconstructed using FastML 3.11 with the options “--seqType NUC --SubMatrix GTR --indelReconstruction ML”.
4. Using the results of this reconstruction, we determined the AT content at the ancestral nodes.

The results of the analysis from section step 2 (see details in the section “Is the AT content of the plastid genome of rhopalocnemis still increasing?” in Results and Discussion) showed that the AT content of rhopalocnemis is likely decreasing. One potential explanation for this result could be that FastML incorrectly reconstructed the ancestral sequences. To test for this, we simulated (see details below) the evolution of a genome similar to that of rhopalocnemis (with the same characteristics of nucleotide substitution and indel processes) but with an AT content that does not change over time. Subsequently, for each of the 1000 simulation replicates, we reconstructed the ancestral sequences using FastML and checked whether FastML would yield the same result (i.e., that the AT content decreases over time). Specifically, the analysis was arranged as follows:

1. The process generating substitutions during evolution with a constant AT content, using the GTR+Gamma model, is defined by the substitution matrix and the alpha parameter of the gamma distribution. Both were determined for the evolution of rhopalocnemis using IQ-TREE 1.6.12 on the basis of the multiple alignment of the plastid genomes of rhopalocnemis. IQ-TREE was instructed to use the substitution model GTR+Gamma with the option “-m GTR+F+G”.
2. Assuming that the lengths of indels are distributed according to the Zipf distribution, the process generating indels is described by two values. The first is the ratio of the rate of indels to the rate of substitutions. The second is the power parameter of the Zipf distribution. Both values were determined by the script lambda.py, included with the DAWG 1.2 program (Cartwright 2005), on the basis of the multiple alignment of the plastid genomes of rhopalocnemis.
3. The sequence simulation was performed using DAWG. The characteristics of the processes generating substitutions and indels, which are required as inputs for DAWG, were calculated as described above. A total of 1000 replicates of the genome evolution simulation were performed.
4. For each of the 1000 replicates, ancestral sequence reconstruction was performed on the sequences resulting from the simulated evolution using FastML with the options “--seqType NUC --SubMatrix GTR --indelReconstruction ML”.
5. AT contents in ancestral nodes were determined on the basis of the results of the reconstruction. A metric characterizing the change in AT content over time, which we call the “AT gradient,” was calculated on the basis of the AT content. The AT gradient is calculated as the ratio of the following two values: a) The sum of changes in AT content across all branches. For example, if on a certain branch, the AT content in the descendant node was 85% and that in the ancestral node was 80%, then the change in AT content was 5%. The value in the ancestral node is subtracted from the value in the descendant node. b) The sum of the lengths of the tree branches.

Thus, the AT gradient is how much the AT content changes during evolution per unit of branch length. In other words, it is how much the AT content changes over the time during which, on average, one substitution occurs at each position. A positive AT gradient indicates an increase in AT content over time, whereas a negative AT gradient indicates a decrease in AT content over time.

To determine whether the decline in the AT content of rhopalocnemis is likely just an error in the reconstruction of ancestral sequences by FastML, we compared the AT gradient obtained from the reconstruction of ancestral sequences based on real modern sequences of rhopalocnemis with the AT gradient obtained from the reconstruction of ancestral sequences based on simulated modern sequences of rhopalocnemis (whose evolution in simulations occurred such that the AT content did not change over time). The comparison was conducted by calculating the percentile that the real AT gradient constitutes compared to 1000 AT gradients obtained in 1000 simulation replicates.

Reconstruction of the history of indels is a challenging problem, and errors in the reconstruction of the history of indels can potentially distort the results of the analysis of AT content evolution. Therefore, the analyses described above were conducted not only for the alignment of plastid genomes as a whole but also separately for the alignment of plastid genomes processed with Gblocks using the options “-t=d -b3=4”. Since Gblocks, when run in this way, removes all columns with gaps, the reconstruction of indel evolution is not required in this case. Supplementary Note 2 provides both the results without Gblocks and the results with Gblocks.

### Other analyses

Codon frequency calculations were performed using the Cusp program from the EMBOSS 6.5.7 software suite.

The phylogenetic trees in this article were drawn using TreeGraph 2.14.0 (Stöver & Müller 2010). A diagram demonstrating the distribution of polymorphism metric values throughout the genome (Figure 2c) was constructed using pyGenomeTracks 3.8 (Lopez-Delisle et al. 2021). The values of the polymorphism metrics were calculated for 100-bps windows of multiple alignment of seven samples of rhopalocnemis. The coordinate of the window was considered the coordinate of the 51st base pair of the window. In cases where the window crossed the edge of the alignment, the remaining part of the window was taken from the opposite end of the alignment because the plastid genomes are circular. The windows were taken with a step of 1 bp.

Analysis of correlations between different metrics of polymorphism (Supplementary Figure 1) was performed for 100-bps windows with a step of 100 bps because the analysis of the statistical significance of the correlation implies independence between the data, meaning that the windows should not overlap.

The structure of *trnE* of *Arabidopsis thaliana* for Figure 4a was determined by tRNAscan-SE and visualized by VARNA (Darty et al. 2009).

## Acknowledgements

We are grateful to Dmitry Lyskov for assistance in collecting the samples.

## Data availability

Sequencing reads are available on the NCBI website as part of BioProject PRJNA495456. The NCBI accession codes of the plastid genome sequences and their annotations can be found in Table 2. The computer code used in this study is available at https://doi.org/10.6084/m9.figshare.30225256. The code used to calculate polymorphism is also available at https://github.com/shelkmike/Polymorphism_suite.

