## Supplementary material for "The plastid genome of the non-photosynthetic plant *Rhopalocnemis phalloides* is one of the most polymorphic genomes known": supplementary_note_1__correction_of_errors_in_previous_rhopalocnemis_assemblies.docx

### Supplementary Note 1. On the correction of errors in previous assemblies of plastid genomes of rhopalocnemis

In this work, we reassembled the plastid genomes of all 7 samples. We found that the new assemblies of the plastid genomes of sample 1387 and sample WS differ from those deposited on the NCBI website (NCBI accession codes MK036331.1 and PQ849602.1, respectively). Coverage analysis (Figures X1-X10) revealed that in the new assemblies, the coverage is more uniform, indicating that they are likely more accurate.

In the new assembly of the plastid genome of sample 1387, compared with the old assembly, there is an insertion of 210 bps in the CDS of the *ycf1* gene. This insertion increases the length of the *ycf1* gene’s CDS from 2265 to 2475 bps. This insertion was made in a region containing multiple copies of tandem repeats—apparently, the number of tandem repeat copies was previously determined incorrectly. Although the insertion is within a CDS, it does not disrupt the open reading frame, as the length of the insertion is a multiple of three and the insertion does not contain stop codons.

In the new assembly of the plastid genome of sample WS, 105 bps in the first (of two) intron of the *clpP* gene were deleted. These base pairs are located in the plastid genome deposited on the NCBI website in the region from 15,765 to 15,869 inclusive. This change reduces the length of the intron from 492 bps to 387 bps. The change involves the deletion of one copy of a tandem repeat with a long monomer.

Below (Figures X1-X9) are diagrams of Illumina read coverage and AT content in 7 samples of rhopalocnemis.

For samples 1387 and WS, coverage diagrams depict both before and after error correction. Since high AT content hinders sequencing, to reduce the impact of AT content on coverage, we applied AT content correction to each sample using the correctGCBias program from the DeepTools suite. The correction was made on the basis of a window size equal to the insert size of the corresponding library. The AT content correction is not perfect; therefore, some coverage fluctuations remain even after it. Additionally, sample preparation for different samples was performed using different methods, which likely resulted in stronger fluctuations in some samples and weaker fluctuations in others.

The coverage in the figures is shown for each position, while AT contents are averaged over a window whose size equals the average insert size, with the center of the window located at the given position.

Since aligning reads to the edges of contigs is challenging, to demonstrate the continuity of coverage along the entire length of circular plastid genomes, in addition to the coverage diagram for plastid genomes in the main orientation, we also created coverage diagrams for genomes where the first 10,000 bps are moved to the end.

To reduce the number of mitochondrial and nuclear Illumina reads aligned to the plastid genome, after aligning the reads, we retained only those with at least 50 bps aligned and a similarity of at least 98% in the aligned part.

For samples MLP and WS, not only Illumina reads but also PacBio HiFi reads (that is, high-accuracy PacBio reads) were available, and for the DWS sample, not only Illumina reads but also PacBio CLR reads (that is, low-accuracy PacBio reads) were available. The alignments of PacBio reads to plastid genomes are shown in Figure X10. In particular, these alignments confirm our correction of the WS genome assembly.

To reduce the number of mitochondrial and nuclear PacBio reads aligned to the plastid genome, after alignment, we retained only those reads with at least 10,000 bps aligned. For the HiFi reads (samples MLP and WS), we required a similarity to the plastid genome sequence in the aligned part of the read of at least 98%, and for the CLR reads (sample DWS), we required at least 90%. Unlike the coverage by Illumina reads, we did not adjust the coverage by PacBio reads for AT content because AT content has little effect on coverage by PacBio reads.

#### Figure X1. Coverage distribution by Illumina reads for sample MLP.

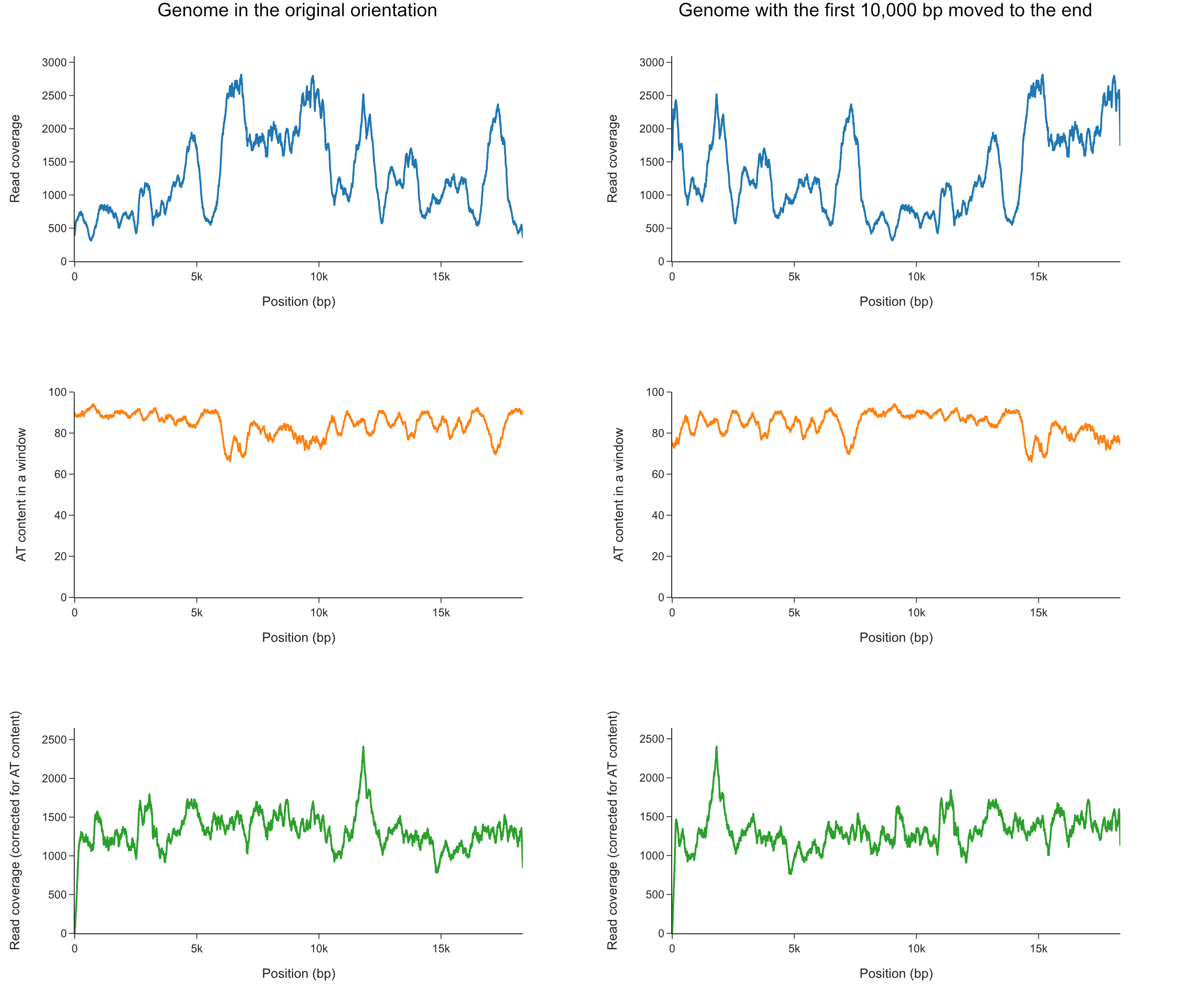

#### Figure X2. Coverage distribution by Illumina reads for sample WS (before fixing the error).

The region with the error is marked by a red arrow in the coverage diagram.

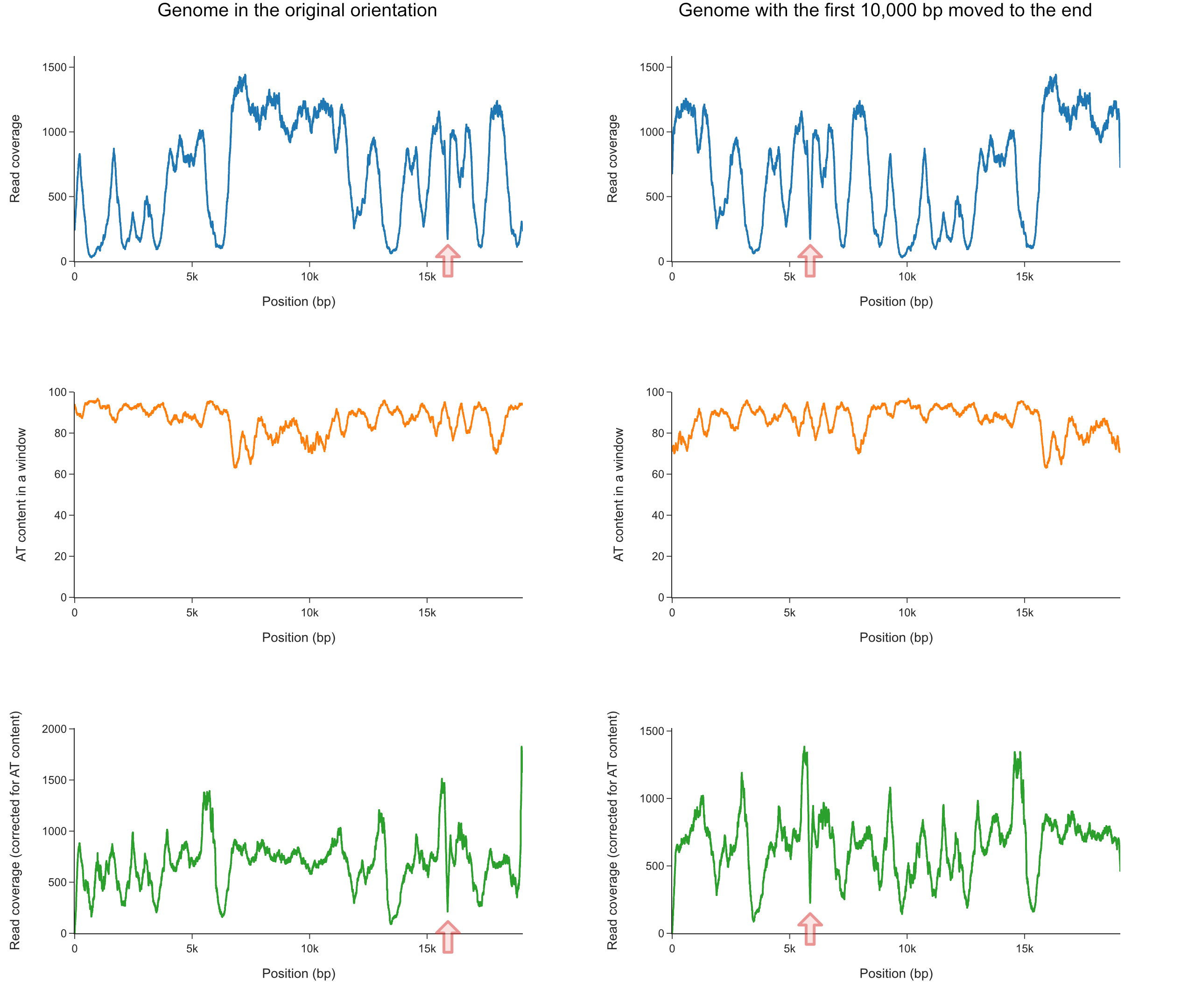

#### Figure X3. Coverage distribution by Illumina reads for sample WS (after fixing the error).

The region where the error was fixed is marked by a red arrow in the coverage diagram. For comparison, see Figure X2.

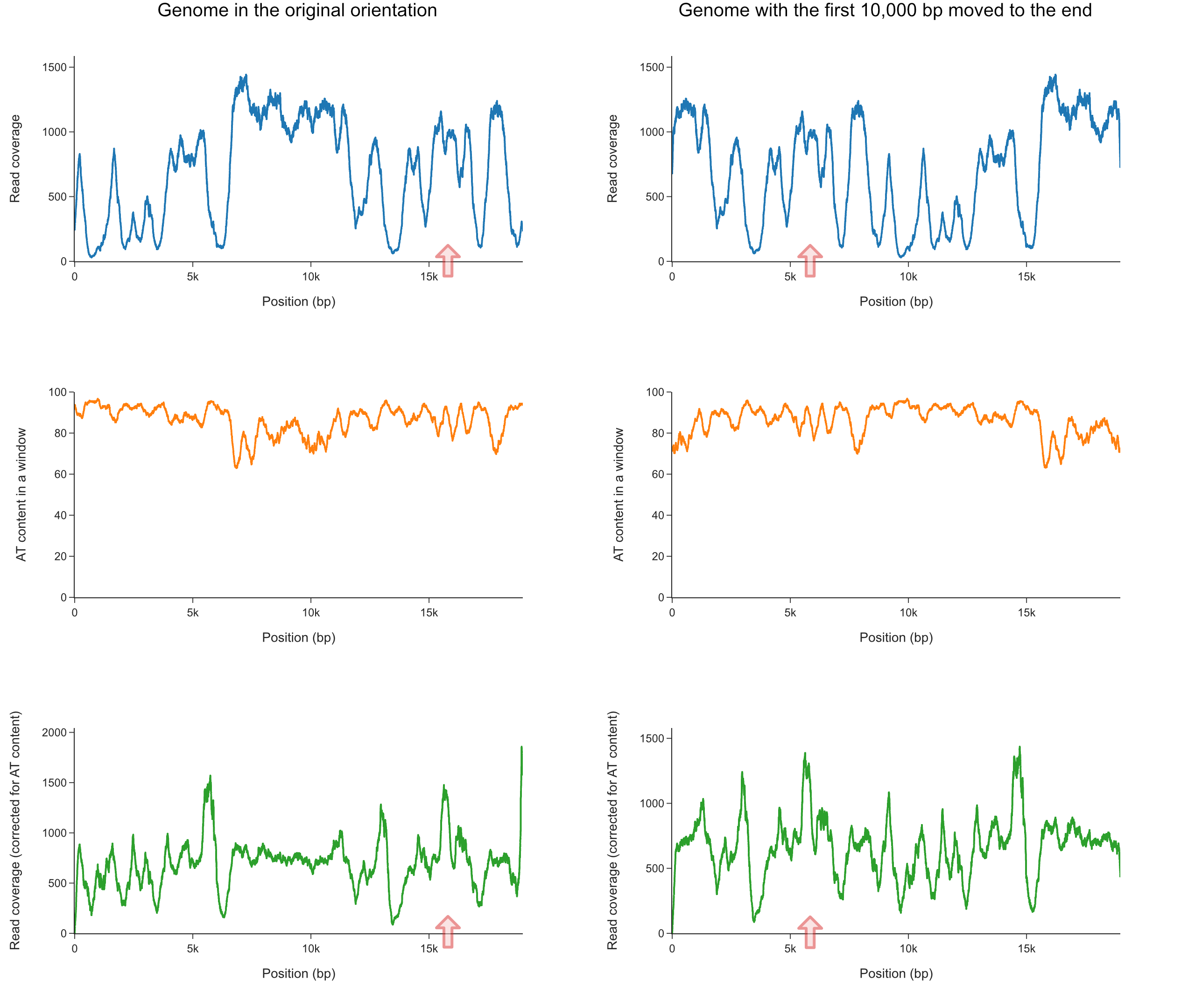

#### Figure X4. Coverage distribution by Illumina reads for sample DWS.

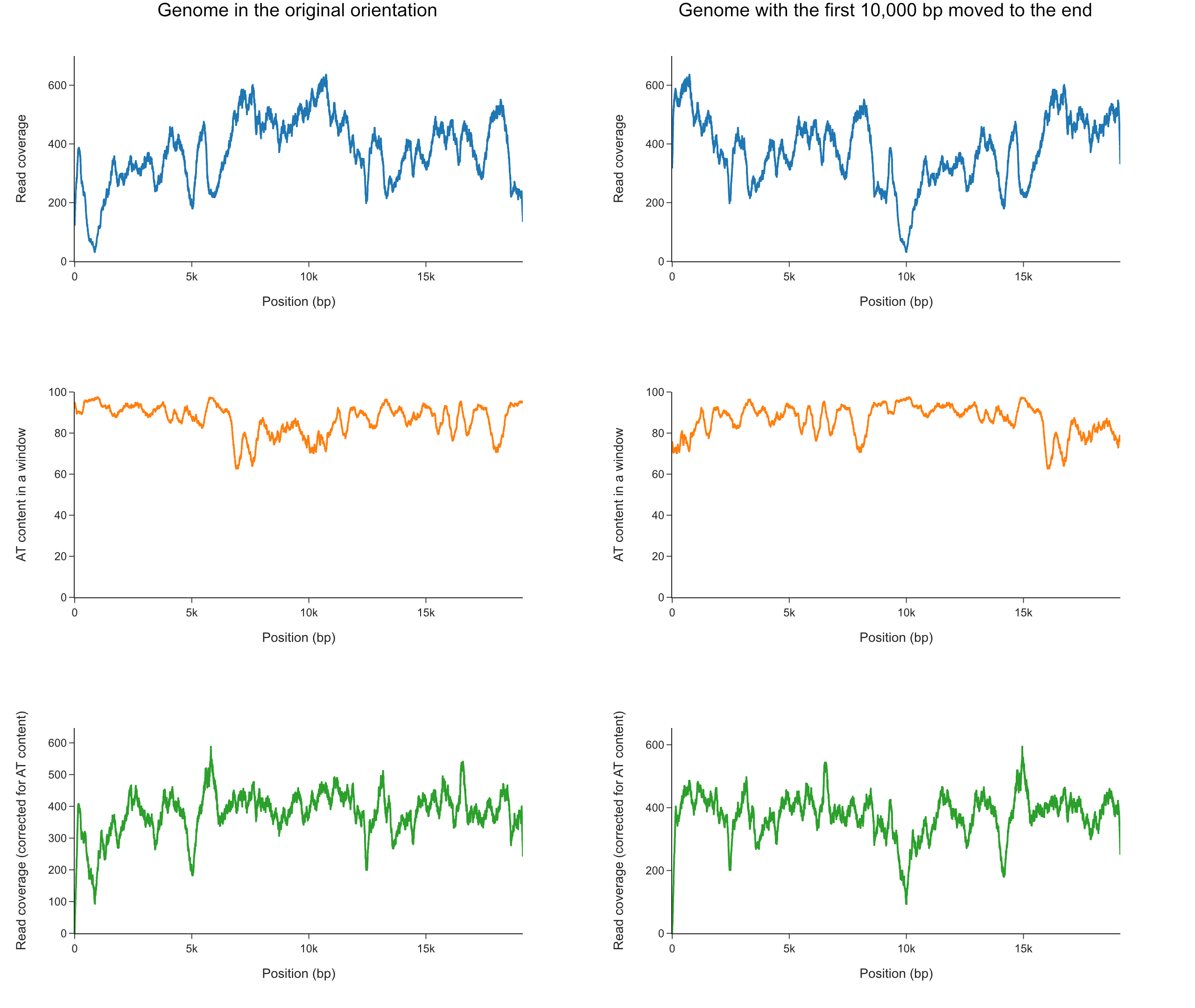

#### Figure X5. Coverage distribution by Illumina reads for sample 2194.

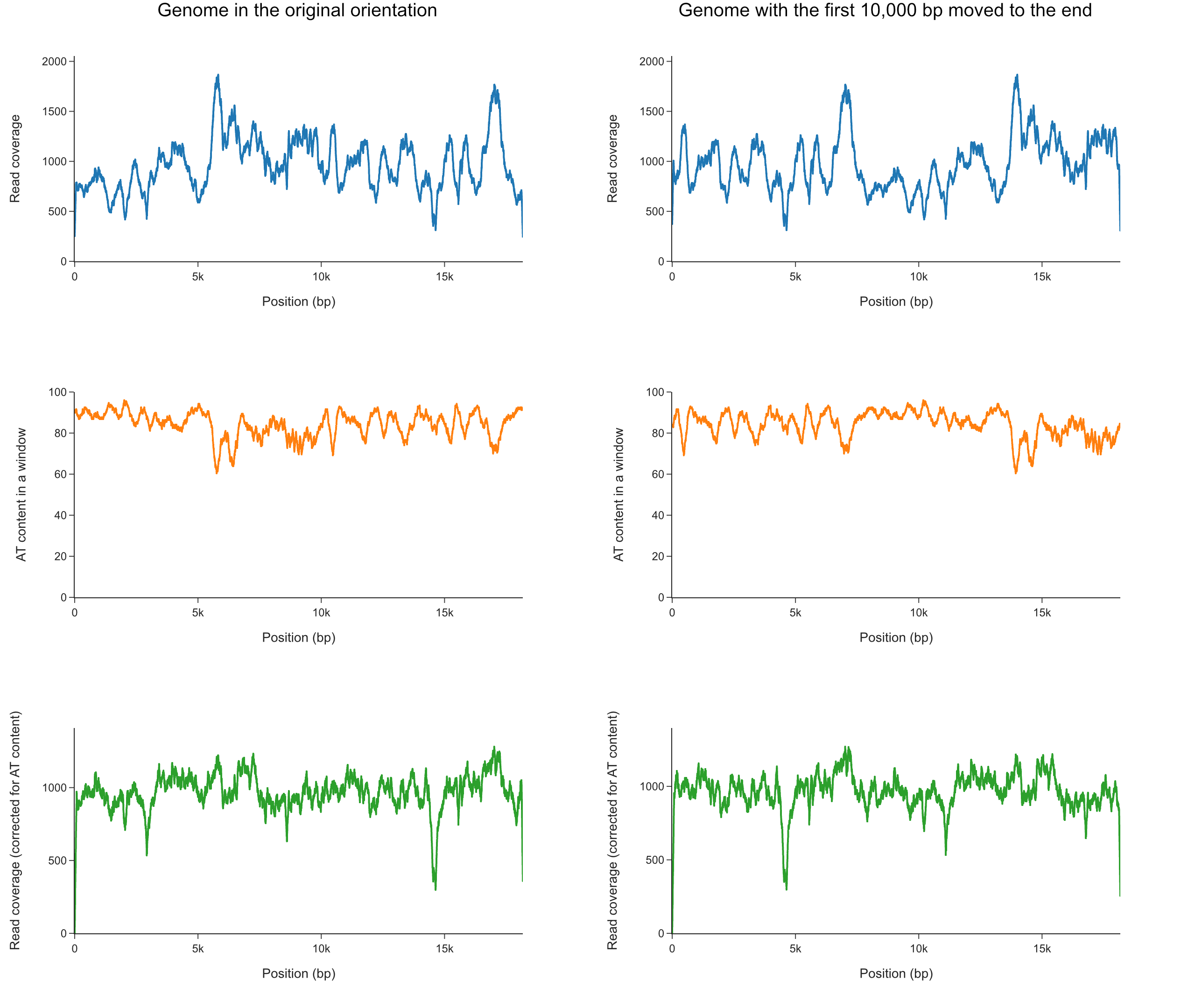

#### Figure X6. Coverage distribution by Illumina reads for sample 3451.

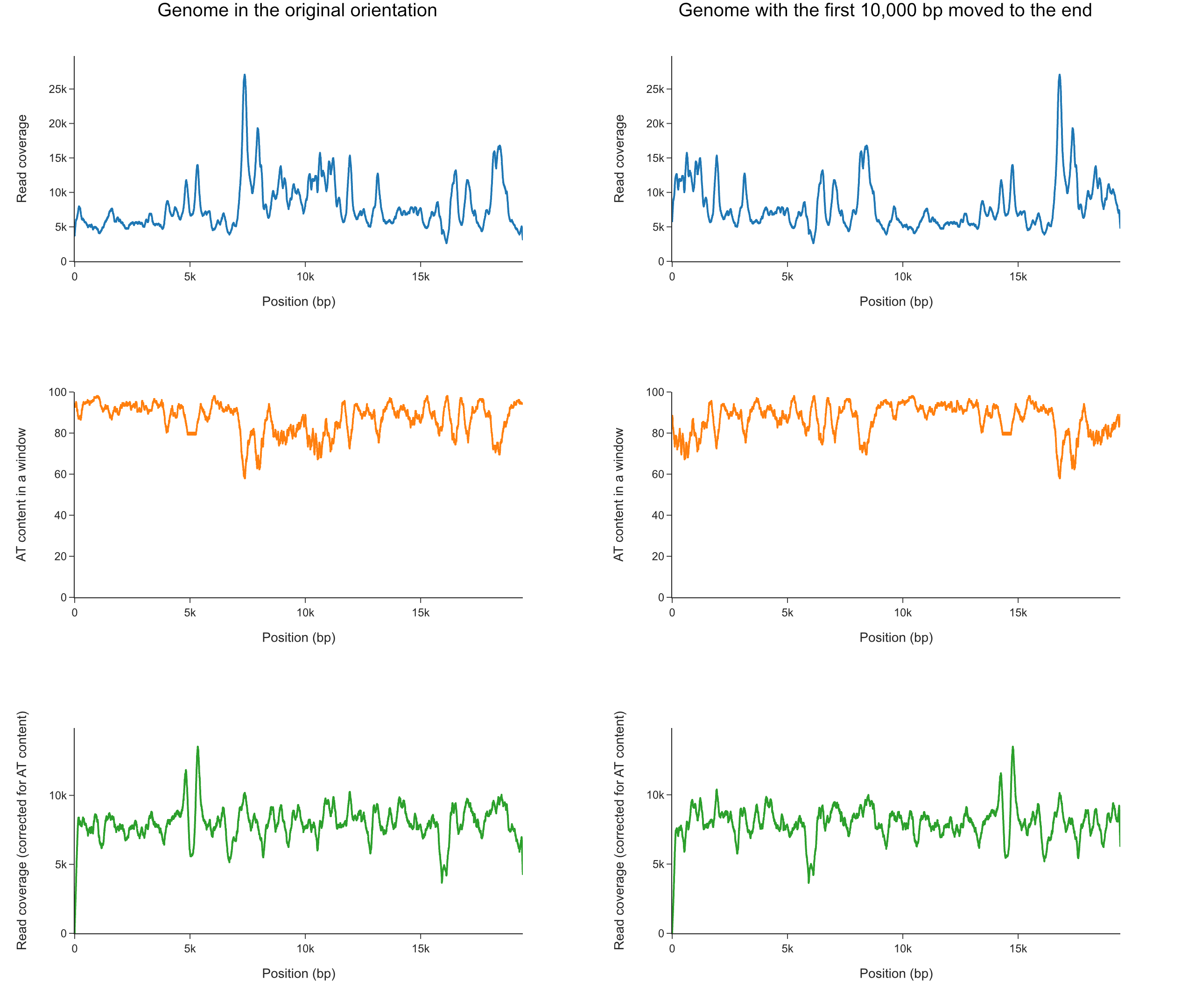

#### Figure X7. Coverage distribution by Illumina reads for sample 1387 (before fixing the error).

The region with the error is marked by a red arrow in the coverage diagram.

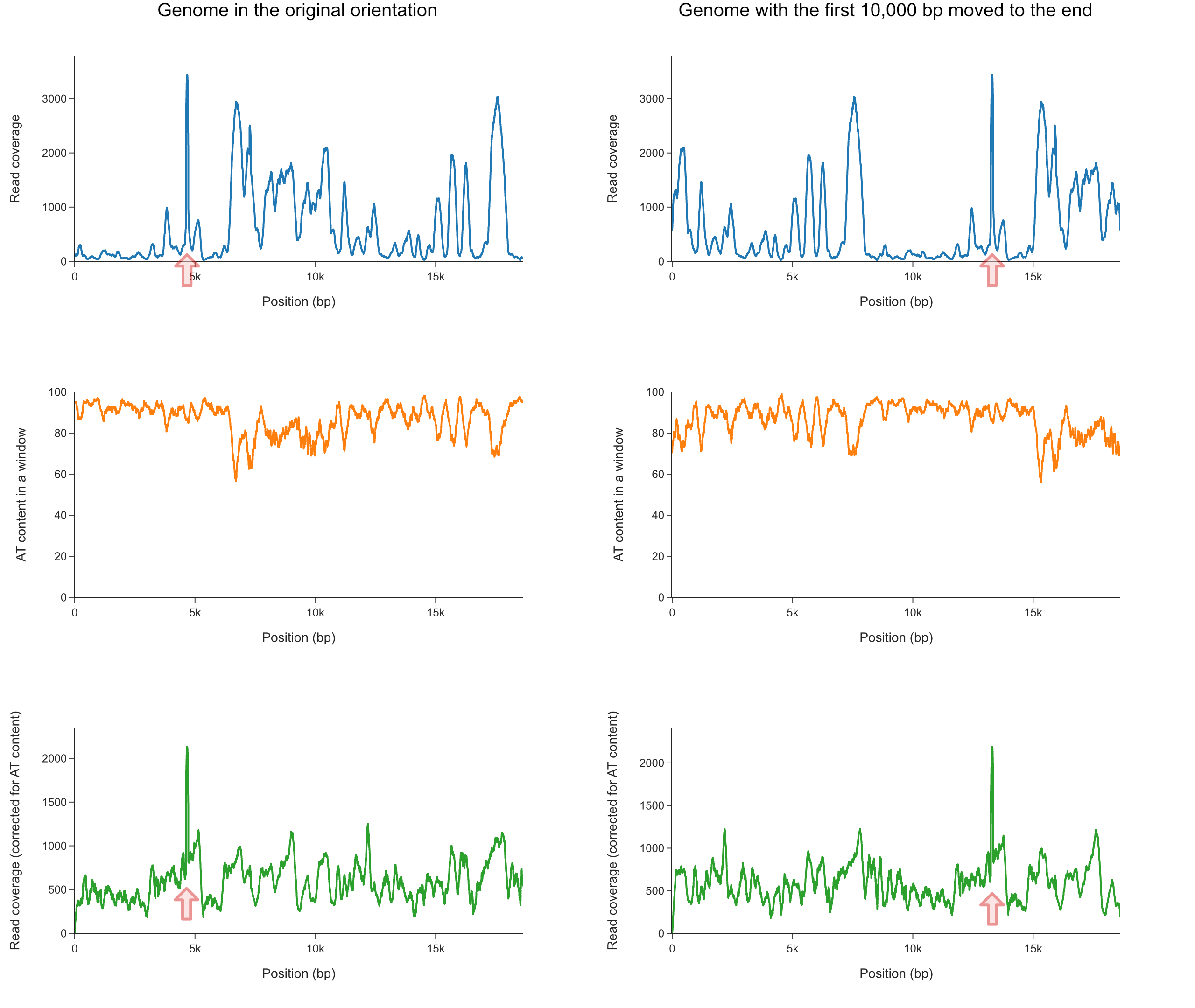

#### Figure X8. Coverage distribution by Illumina reads for sample 1387 (after fixing the error).

The region where the error was fixed is marked by a red arrow in the coverage diagram. The absence of a fluctuation in the coverage is noticeable after a correction for AT content. For comparison, see Figure X7.

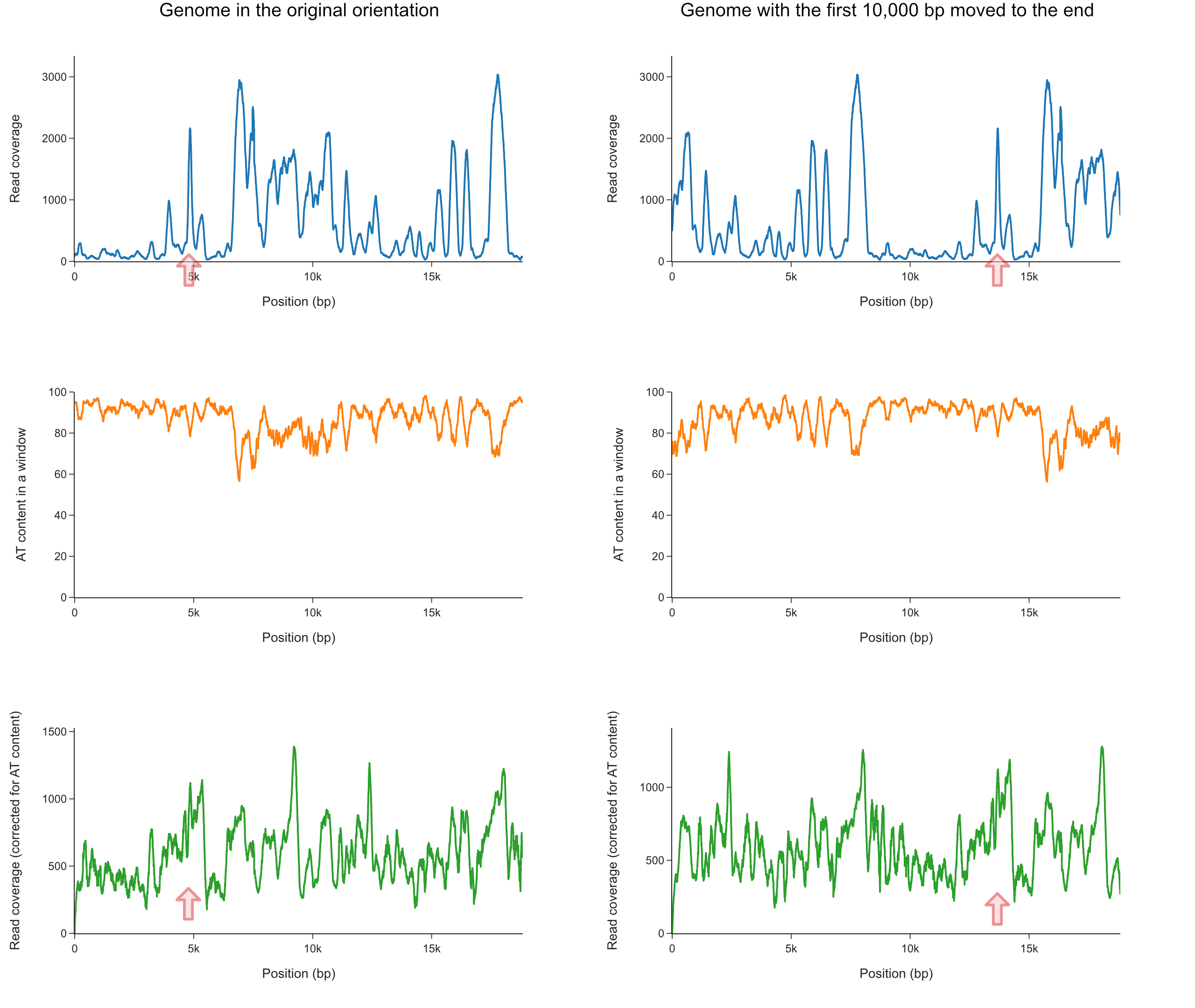

#### Figure X9. Coverage distribution by Illumina reads for sample 1002.

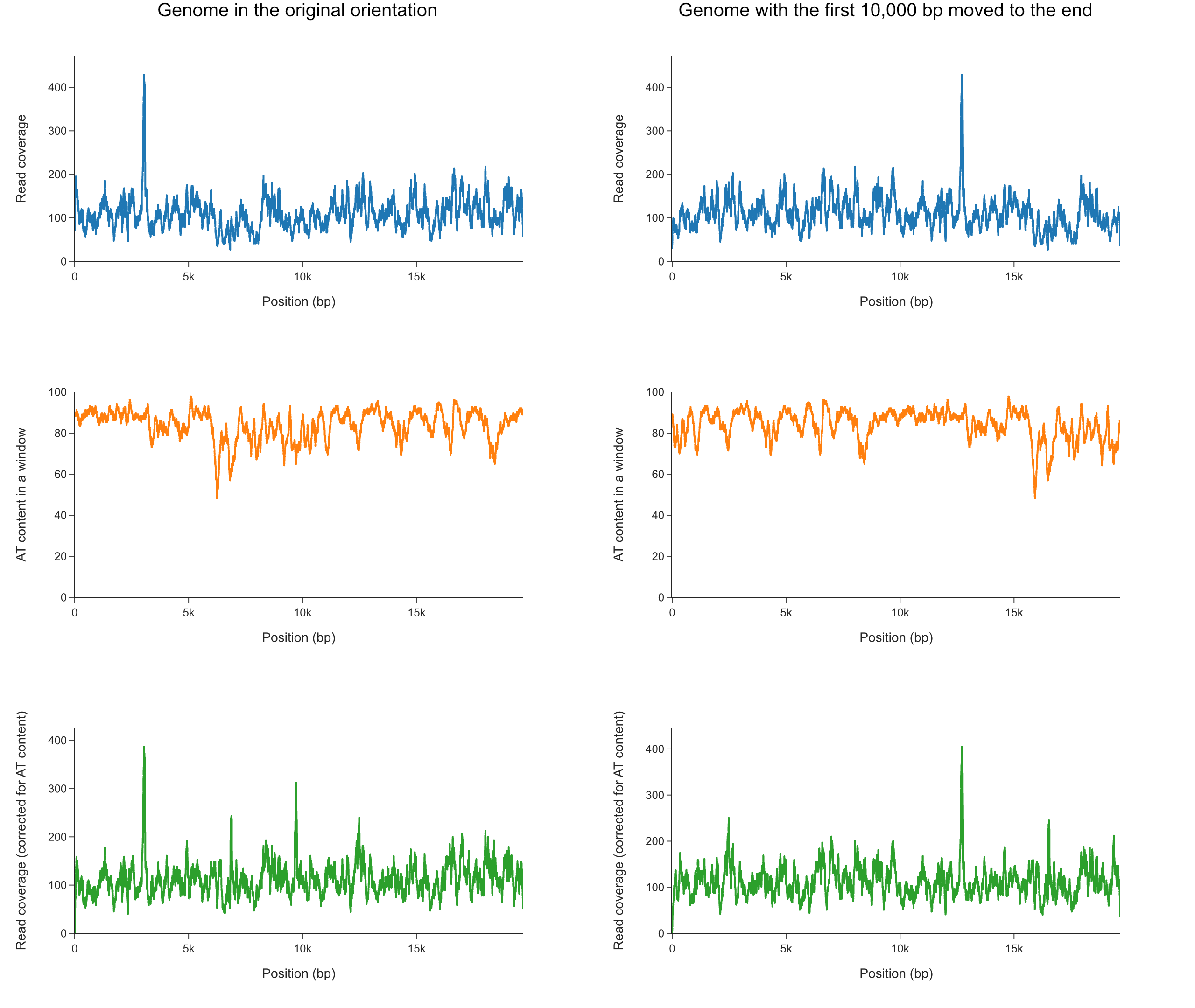

#### Figure X10. Coverage distribution by PacBio reads for samples MLP, WS, DWS.

The coverage gradually decreases toward the ends because of the requirement that at least 10,000 bp of a read should align in one region; this causes underrepresentation of reads that span the contigs’ edges. The region with the error in sample WS is marked by a red arrow.

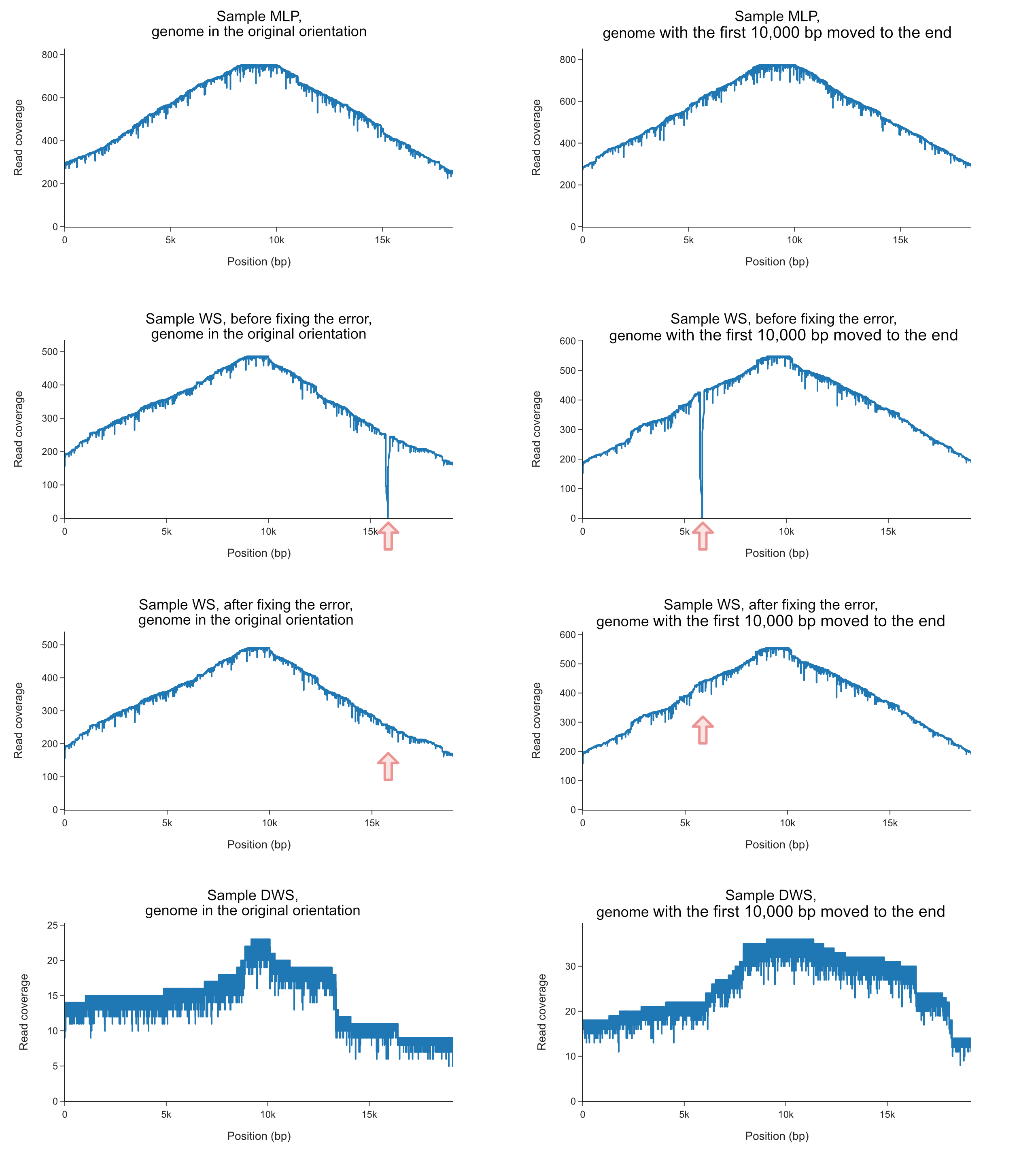
