## Supplementary material for "The plastid genome of the non-photosynthetic plant *Rhopalocnemis phalloides* is one of the most polymorphic genomes known": supplementary_note_2__additional_information_on_AT_content_evolution.docx

### Supplementary Note 2. Additional information on the analysis of AT content evolution

The results of the reconstruction of the ancestral AT content in the plastid genomes of rhopalocnemis are presented in Figure Y1a. The value is not provided for the common ancestor of all seven samples because the analysis was conducted only on the sequences of the plastid genomes of rhopalocnemis, which does not allow the point on the ancestral branch where the tree should be rooted to be precisely determined in this analysis.

It is evident that all ancestral AT contents are even higher than the AT contents of modern plastid genomes of rhopalocnemis.

To verify that this was not an artifact of reconstruction, we simulated evolution with a constant AT content of genomes that are as AT-rich as the plastid genomes of rhopalocnemis and have the same processes generating substitutions and indels during evolution (for details, see the "Materials and Methods" section in the main text of the article). A total of 1000 simulation replicates were generated, and then ancestral sequences were reconstructed for each replicate in the same way as for the real sequences of rhopalocnemis. The change in AT content on the tree was assessed by a measure that we call the AT gradient; it is positive when, overall, the AT content increases across the tree and negative when it decreases. In the ancestral reconstruction made from the leaves obtained in all 1000 simulations, the AT gradient was higher than that of the real rhopalocnemis, which suggests that the decrease in AT content we observed was not the result of an error in ancestral sequence reconstruction.

**Figure Y1. Analysis of ancestral AT contents in plastid genomes of rhopalocnemis without Gblocks.** (a) AT contents in modern sequences of rhopalocnemis as well as in reconstructed ancestral sequences of rhopalocnemis. The scale under the tree indicates the rate of substitution. (b) A sina plot with the distribution of AT gradients in simulated evolution with constant AT contents (each gray dot is a replicate), as well as the AT gradient resulting from the analysis of the evolution of real rhopalocnemis sequences (i.e., based on the values from (a); red dot).

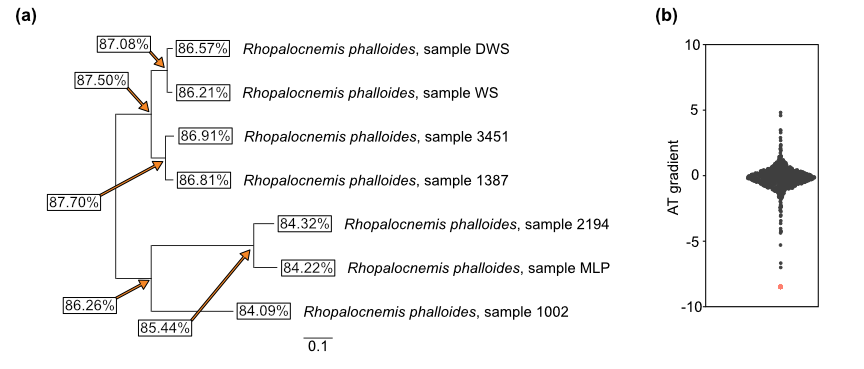

It is known that correct reconstruction of ancestral indels is more complex than reconstruction of ancestral substitutions. To eliminate the possible influence of erroneous reconstruction of ancestral indels, we also conducted an analysis similar to the one described above but on an alignment in which all columns with at least one gap were removed by Gblocks. Additionally, Gblocks removed sections in the alignment that its algorithm considered possibly misaligned. The results of this analysis are presented in Figure Y2. As with the analysis without Gblocks, the reconstruction of ancestral sequences revealed that the AT content of rhopalocnemis decreased. Interestingly (Figure Y2b), when reconstructing ancestral sequences from leaves obtained through simulations with unchanged AT content for genomes without indels it turns out that the AT content slightly decreased rather than remaining unchanged, although the decrease in AT content of the real rhopalocnemis was stronger than that in all 1000 simulations. The reason for this is unknown to us.

Due to the complexity of reconstructing ancestral sequences of very rapidly mutating genomes, we cannot rule out that our result (that the AT content in rhopalocnemis decreases) is still an artifact.

**Figure Y2. Analysis of ancestral AT contents in plastid genomes of rhopalocnemis** **after processing multiple alignments with Gblocks.** (a) AT contents in modern sequences of rhopalocnemis as well as in reconstructed ancestral sequences of rhopalocnemis. The scale under the tree indicates the rate of substitution. (b) A sina plot with the distribution of AT gradients in simulated evolution with constant AT contents (each gray dot is a replicate), as well as the AT gradient resulting from the analysis of the evolution of real rhopalocnemis sequences (i.e., based on the values from (a); red dot).

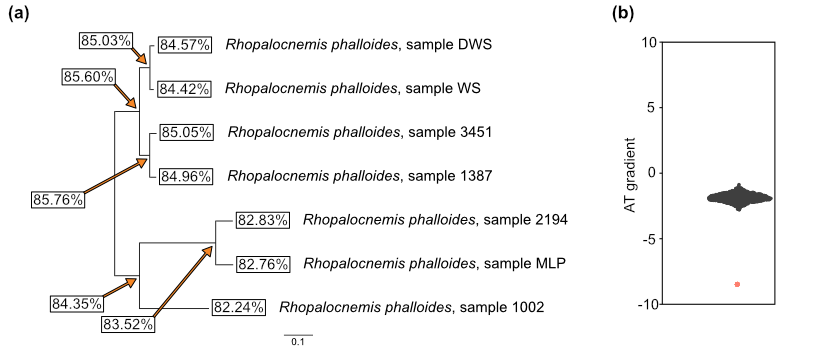
